# Sex-Dependent Epigenomic and Transcriptomic Reprogramming Links Maternal Obesity to Cardiac Remodeling in Adult Offspring

**DOI:** 10.1101/2025.04.15.648971

**Authors:** Tim D. Wilson, Elysse A Philips, Yem J Alharithi, Cameron Broberg, Joshua Vanderpool, Shauna Rakshe, Suzanne S. Fei, Brett A. Davis, Sheryl Koch, Zhehao Zhu, Lyndsey E. Shorey-Kendrick, Sushil Kumar, Lucia Carbone, Yabing Chen, Jamie O. Lo, Jack Rubinstein, Susan B. Gurley, Karina Nakayama, Sandra Rugonyi, Alina Maloyan

**Author notes:** **Corresponding Author:** Alina Maloyan, PhD, FAHA. Knight Cardiovascular Institute, Oregon Health and Science University, 3181 SW Sam Jackson Park Rd, Portland, Oregon, 97239.

## Abstract

Maternal obesity during pregnancy predisposes the offspring to a high risk of developing cardiovascular and metabolic diseases later in life. This study investigated cardiac perturbations caused by maternal obesity by utilizing a mouse model of maternal high-fat diet (HFD)–induced obesity that recapitulates metabolic abnormalities observed in humans. Our study revealed that offspring of HFD-fed mothers (Off-HFD) exhibit a progression of obesity, dyslipidemia, and metabolic inflexibility when compared with offspring of regular diet–fed mothers (Off-RD). Deeper investigation of cardiac function revealed profound functional, metabolic, vascular, and immune perturbations in adult Off-HFD mice, with marked sex-specific differences. Although both male and female Off-HFD mice developed progressive cardiac hypertrophy, male offspring exhibited a more severe phenotype characterized by hypertension, increased vascular stiffness, impaired cardiac function, and myocardial fibrosis. To identify potential mechanisms underlying these changes, we performed DNA methylation analysis in collected hearts of newly weaned and adult offspring. This analysis revealed extensive, sex-dependent alterations in DNA methylation within or nearby genes involved in cardiac development, lipid metabolism, hypertrophic growth, and inflammatory signaling. Importantly, many of these epigenetic alterations persisted into adulthood, suggesting that maternal obesity establishes a durable molecular memory in the offspring heart. Consistent with these findings, transcriptome analysis of adult hearts revealed activation of gene programs associated with heart failure and pathological cardiac remodeling in male Off-HFD mice, whereas female Off-HFD mice showed activation of pathways consistent with adaptive or cardioprotective responses. Together, these findings demonstrate that maternal high fat diet induces early-life epigenetic remodeling in the offspring heart that persists into adulthood and is associated with sex-specific metabolic, functional, vascular, and immune dysregulations. By linking early epigenomic changes to adult cardiac disease susceptibility, this study identifies potential developmental windows for preventive and early therapeutic interventions aimed at reducing cardiovascular risk in offspring exposed to maternal obesity *in utero*.

## INTRODUCTION

Obesity has become a rapidly escalating public health concern. Obesity has become a rapidly escalating public health concern. Over the past 30 years, the prevalence of obesity, defined as a body mass index (BMI) of over 30, in the United States has doubled (1). Globally, in 2025, 17% of adults—approximately 890 million people—were living with obesity, while 43%—approximately 2.5 billion people—were overweight (BMI of 25-29.9) (2). A similar trend has been observed among women of reproductive age (15–49 years) (3) and as of 2020, more than 40% of women aged 20–39 years are estimated to be living with obesity (4), underscoring the growing magnitude of this public health issue.

A successful pregnancy depends on the tightly regulated transfer of nutrients to the fetus, a process governed by extensive maternal–fetal communication. Consequently, alterations in maternal health and disruption of this communication can leave long-lasting imprints on the fetus, a phenomenon known as developmental programming. These changes may result in structural and physiological abnormalities that manifest as disease later in the offspring’s life. Epidemiological evidence links obesity during pregnancy to increased risk of neurodevelopmental disorders, including cerebral palsy, attention-deficit disorder, cognitive delay, and autism spectrum disorder (5–7), as well as major congenital anomalies, such as neural tube defects (8). Similarly, epidemiological and clinical studies provide consistent evidence that maternal obesity is associated with cardiovascular dysfunction in offspring across the life course (9–12). Indeed, maternal obesity is now increasingly recognized as a significant risk factor for cardiovascular dysfunction in offspring during childhood and early adulthood (13). For example, an epidemiological study of 37,709 individuals with birth records in the United Kingdom (UK) from 1950 to 2013 found offspring of mothers with an elevated BMI to have significantly higher rates of hospital admission for cardiovascular events and premature death than offspring of mothers with normal BMI, even after adjustment for maternal age, socioeconomic status, and offspring sex (14). Consistent with these findings, another UK study of 1,555 mothers with obesity showed that, compared with children born to women with a normal BMI, children born to mothers with obesity exhibit evidence of cardiac remodeling, such as increased interventricular septal thickness, posterior wall thickness, and relative wall thickness after adjustment for confounding factors (10).

Large population-based studies further support a dose-dependent relationship between maternal obesity and offspring cardiovascular risk. A Swedish cohort study that followed more than 2.3 million infants born between 1992 and 2016 demonstrated offspring cardiovascular disease risk tended to increase in conjunction with the severity of maternal obesity (13). In the same study, sibling-cohort analysis also showed a positive trend between maternal BMI and offspring cardiovascular disease rates, supporting a potential intrauterine contribution to this association (13). Evidence from systematic analyses is consistent with these observations: a literature search of 1,589 publications identified reduced systolic function and increased interventricular septal thickness in children born to women with obesity (15). Additional prospective and smaller-cohort studies have linked maternal obesity with adverse cardiometabolic and vascular outcomes in offspring. In a prospective study of 1,400 young adults born in Jerusalem, high maternal BMI was associated with offspring dyslipidemia and higher systolic and diastolic blood pressure (16). Similarly, a Belgian study of 240 mother–child dyads found a positive association between maternal pre-pregnancy BMI and increased mean arterial pressure in early life, independent of the child’s current BMI (17). Together, these findings indicate that maternal obesity creates an adverse early-life environment that may program lifelong susceptibility to cardiovascular disease in offspring. However, the mechanisms underlying this increased risk remain poorly understood.

We have developed a mouse model of maternal high-fat diet–induced obesity that reflects key metabolic abnormalities seen in offspring of mothers with obesity including obesity, glucose intolerance, hyperinsulinemia and asthma (18–21). Using this model, we provide insight into how maternal obesity leads to long-term cardiac dysfunction in offspring. By combining physiological, histological, epigenomic, and transcriptomic analyses, we identified persistent sex-specific molecular changes linked to changes in lipid handling and utilization, metabolic flexibility and extracellular matrix remodeling, and altered regulatory pathways.

## MATERIALS AND METHODS

### Study approval

All animal experiments were approved by Oregon Health & Science University’s Institutional Animal Use Committee (IACUC protocol # IP00432).

### Antibodies

Total OXPHOS Rodent Western Blot Antibody Cocktail was purchased from Abcam (Waltham, MA, cat # ab110413). Antibodies against Vinculin, p62, Rubicon, ATG 7, and LC3-II were purchased from Cell Signaling (Danvers, MA, cat # 4650, 5114, 8465, 8558, and 12741 respectively). Antibody against the autophagy regulator TFEB was purchased from Invitrogen (Waltham, MA, cat #PA1-31552).

### Mouse model of maternal obesity

All mouse studies were done using wild-type FVB/N mice. Mice were kept under a 12-hour light/dark cycle in temperatures between 18–23 degrees Celsius with 40–60% humidity, in stress-free conditions. Mice were caged in groups of three to five whenever possible. Food and water were given ad libitum, and body weight was determined weekly. To induce maternal obesity, a high-fat diet (HFD, Teklad Cat#TD.06415) or its control, a regular diet (RD, PicoLab® Laboratory Rodent Diet Cat # 5L0D), was given to virgin female FVB/NJ mice from six weeks of age and throughout the entire study. The diet compositions have been reported before (18–21). After eight weeks of dietary intervention, RD- and HFD-fed female mice were bred to an age-matched RD-fed male. All offspring were fed the RD only starting from weaning and throughout their life.

**Body fat percentage** was assessed using an EchoMRI Analyzer E26-206-RMT (EchoMRI LLC, Houston, TX) as previously described (18). All outcomes were normalized to body mass at the corresponding time point.

### Metabolic phenotyping

Oxygen consumption and respiratory exchange ratio (RER) were measured in individually-housed mice by Vanderbilt University’s Mouse Metabolic Phenotyping Core using the Promethion from Sable Systems (Las Vegas, NV).

**Mean arterial pressure** was non-invasively measured using a CODA® tail-cuff blood pressure system (Kent Scientific, Torrington, CT) after two weeks of training.

**Blood lipid profiling** was performed at the Mouse Metabolic Phenotyping Center of the University of Cincinnati. Plasma triglycerides were measured using the Randox Triglyceride Assay Kit (Randox Laboratories, Kearneysville, WV). Plasma cholesterol levels were determined using the Infinity Total Cholesterol Assay Kit (Fisher Scientific, Waltham, MA). Plasma phospholipids were assessed using the Phospholipids C Assay (Wako Life Sciences, Inc., Mountain View, CA). Plasma non-esterified fatty acids were measured using the Wako HR Series NEFA-HR (2) (Wako Life Sciences, Inc., Mountain View, CA). All measurements were conducted according to the manufacturer’s instructions.

### Pulse wave velocity calculations

Mice were anesthetized with isoflurane using an induction chamber connected with a scavenger canister. During the examination, the animals were maintained under gaseous anesthesia through a nose cone (1.5–2% isoflurane in 0.4 L/min in pure oxygen) and heart rate, respiration frequency, and body temperature were monitored. The abdomen was shaved using depilatory cream (Nair, Church & Dwight Canada Corp., Mississauga, ON, Canada) and coated with Aquasonic Clear Ultrasound Gel (PLI 03-08, Parker Laboratories, NJ). A 4-mm field-of-view ultrasound probe (MS250, FUJIFILM VisualSonics Inc., Toronto, Canada) held in position by a mechanical arm was used for the acquisitions. B-mode images were obtained with the probe placed above the abdominal aorta to obtain longitudinal images of the vessel with the region of interest located in the focal zone of the UHF46x (46–20 MHz range).

**Photoacoustic imaging** was performed using the Vevo LAZR X20 imaging system equipped with a 21 MHz transducer containing built-in fiber optics (Vevo® LAZR, VisualSonics Inc., Toronto). Before imaging, animals were depilated, anesthetized with 2.5% isoflurane, and positioned on the imaging platform. Average oxygen saturation (sO_2_Av) and total hemoglobin concentration (HbT) were quantified by manually tracing a region of interest over the heart on the ultrasound images using Vevo LAB software. Cardiac lipid accumulation was assessed using the ultrasound platform (Vevo F2, FUJIFILM VisualSonics) equipped with a 40 MHz center-frequency transducer to acquire *in vivo* long-axis B-mode and photoacoustic images. A Nd:YAG pulsed optical parametric oscillator laser capable of generating light from 670 to 2300 nm was used to deliver 5 ns pulses at a repetition rate of 10 Hz. Grayscale B-mode images were used to guide the drawing region of interest within the myocardium, and sO_2_Av and HbT were measured within the region of interest. Wavelengths used were 1210 nm for lipids, 750 nm for oxyhemoglobin, and 850nm for deoxyhemoglobin. Color maps for lipids, HbT, and sO_2_% were generated using a color lookup table and overlaid onto the corresponding B-mode images.

### Transmission electron microscopy

Hearts were perfused with 3.5% glutaraldehyde in 0.1 M/L phosphate buffer, pH 7.4 and embedded in Embed 812 resin (Electron Microscopy Science, Morgantown, PA) for sectioning. The ultrathin sections were visualized by the Multiscale Microscopy Core at OHSU using a FEI Tecnai™ with iCORR™ transmission electron microscope (Thermo Fisher Scientific, Hillsboro, Oregon).

### Protein isolation

Protein was isolated from mouse hearts and lysed using ice-cold radioimmunoprecipitation assay buffer with freshly added protease and phosphatase inhibitors. The resulting mixture was transferred to a 1.5 ml tube and centrifuged at 1000 x *g* for ten minutes at 4 °C to remove cellular debris. The supernatant was then transferred to a fresh tube and the protein concentration quantified using a Pierce Bicinchoninic Acid Protein Assay Kit (Thermo Fisher Scientific, Waltham, MA).

### RNA and DNA isolation

For each sample, RNA and DNA were co-isolated using the AllPrep DNA/RNA Mini Kit (Qiagen, Germantown, MD; cat. no. 80204). Samples were then shipped to Novogene (Sacramento, CA) for reduced representation bisulfite sequencing (RRBS) and RNA-sequencing (RNA-seq) analyses.

### Micro-CT

Fat and lean tissue were visualized in the subcutaneous and visceral space using a Bruker Sky Scan 1276 MicroCT scanner. Briefly, six-month-old mice were anesthetized with isoflurane and scanned for a duration of approximately four minutes with a radiation dose of less than 500 mGy. The electron beam was set to 45 volts and 133 uA, with an exposure time of 217 ms per image through a 0.5 mm aluminum filter. Mice were scanned at a 1024×1024 resolution for a pixel size of 36.4 um, with two-frame averaging and 360-degree scanning. Scans were reconstructed from the proximal end of L1 to the distal end of L5 between −1000 and 1000 Hounsfield units, and a Gaussian filter (level 3) was employed for the clear visualization of soft tissue (22). Regions of interest were drawn manually around the abdominal wall to separate subcutaneous and visceral space, and fat and lean tissue were distinguished by the lowest point of the bi-modal density peaks. Representative images were taken from a reconstruction of the entire scan in CTVox, a 3D volume rendering software from Bruker for visualizing micro-CT datasets, sliced at the level of the sternum and with application of a transfer function to color bone white, fat yellow, and lean tissue red, as determined by the bimodal density peaks.

### Echocardiography

Mice were anaesthetized with isoflurane and subjected to echocardiography at the Small Animal Research Imaging Center at OHSU using Vevo 2100 system (FUJIFILM VisualSonics, Inc, Toronto, ON, Canada). Cardiac function and structure values were obtained from B-mode images from the parasternal long axis (PSLAX). Left ventricular diameter (LVID), posterior wall (LVPW), and intraventricular septum (IVS) were measured in both systole (s) and diastole (d). B-mode PSLAX images were also used to calculate ejection fraction, left ventricular mass, and stroke volume. Stroke volume was also determined as the product of the LV outflow tract cross-sectional area and velocity-time integral on angle-corrected pulsed-wave Doppler to ensure consistency of measurements. LV end-diastolic and end-systolic volumes were calculated from short-axis linear dimensions and long-axis dimensions using established geometric assumptions.

### Western blotting

Myocardial proteins (10–25 µg) were separated on a 4-20% SDS-PAGE gel, then transferred to a polyvinylidene difluoride membrane and blocked for one hour in 5% (w/v) milk in TBS solution with 0.1% Tween 20. Membranes were subsequently incubated overnight with primary antibodies, washed, blocked in 1% milk, and probed with secondary antibodies conjugated to horseradish peroxidase. Western blot membranes were visualized using G:Box and analyzed using GeneTools, both utilities from Syngene (Frederick, MD). All samples were normalized to housekeeping genes.

### Histology

Masson trichrome staining was performed at the OHSU Histopathology Core. Trichrome-stained sections were used to evaluate myocyte cross-sectional area with the aid of ImageJ (National Institutes of Health, Bethesda, Maryland).

### Flow cytometry

The list of antibodies is presented in **Table S1.** In preparation for flow cytometry, nearly 10^6^ cells from individual hearts were first incubated on ice for 30 min in a solution consisting of 1:10 Fc Receptor Binding Inhibitor (eBioscience, Thermo Fisher Scientific) and 1:500 Live/Dead Aqua stain (Invitrogen) in PBS. Cells were then combined with fluorescently labeled monoclonal antibodies as previously described (23) in a solution of PBS + 5% FCS + 1.0 mM EDTA (FACS buffer). After another 30 min incubation on ice, the cells were washed 1x with FACS buffer. Next, the stained cells were treated with permeabilization/fixation buffer (eBioscience) on ice for ten minutes, then washed 1x using permeabilization buffer (eBioscience). The permeabilized cells were then subjected to intracellular staining by incubation with fluorescently labeled monoclonal antibodies in FACS buffer for 30 minutes on ice. Finally, flow cytometry data were acquired on a Cytek Aurora (Cytek Biosciences, Fremont, CA) spectral flow cytometer and analyzed using FlowJo software v9.5 (Ashland, OR).

### RRBS method and analysis

RRBS analysis of the two age groups was performed independently by different groups, therefore methods differ slightly between analyses. Details of RRBS analysis are presented in the Supplemental Materials.

### RNA-sequencing analysis

A detailed description of the RNA-sequencing analysis is provided in the Supplemental Materials.

**Gene Ontology and pathway analysis** is described in the Supplemental Materials.

### Statistical analysis

Means were compared using two-way analysis of variance (two-way ANOVA) followed by Student’s *t*-test (corrected for multiple comparisons) or its non-parametrical equivalent. The obtained *p*-values are indicated in each respective graph. Significance was set at an alpha of 0.05. When no sex-dependent differences were found, the results from males and females were pooled.

## RESULTS

### Physiological changes in mouse offspring of obese mothers

The experimental model is illustrated in **Fig. 1A**. Briefly, six-week-old female mice were maintained on either a regular diet (RD; 13% kcal from fat) or a high-fat diet (HFD; 45% kcal from fat). Following eight weeks of dietary exposure, females were mated with RD-fed males. Offspring born to HFD-fed dams (Off-HFD) exhibited significantly lower birthweights than offspring born to RD-fed dams (Off-RD) (**Fig. 1B**). However, despite the lower birthweight, both male (**Fig. 1C**) and female (**Fig. 1D**) Off-HFD offspring underwent catch-up growth by weaning (three weeks of age) and remained heavier than Off-RD controls throughout their lifespan.

**Figure 1.**
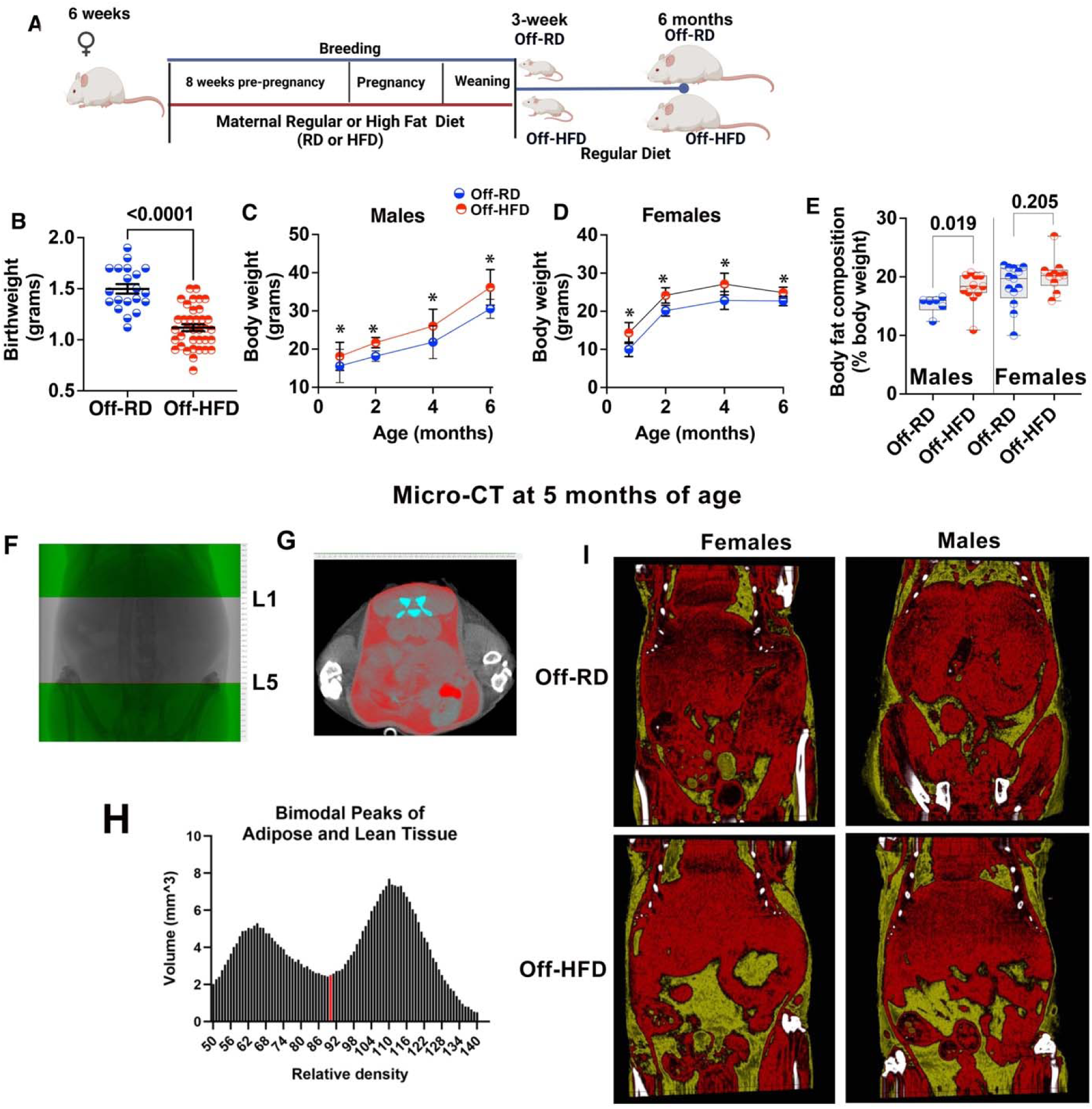
Birthweight and age-dependent changes in body weight and body composition in Off-RD and Off-HFD mice. **A,** Schematic of the mouse model of maternal diet-induced obesity. **B,** Birthweights of Off-RD and Off-HFD. Data shown as individual values; *p*-values are indicated. **C-D,** Body weight trajectories from weaning to 6 months of age in male (**C**) and female (**D**) Off-RD and Off-HFD. Data were analyzed by two-way ANOVA followed by multiple unpaired *t*-tests. N=20–44 per maternal diet/age group; *p* < 0.05 for Off-HFD vs. Off-RD at each age. **E,** EchoMRI analysis showing increased adiposity in adult male, but not female, Off-HFD mice compared with Off-RD controls. *P*-values are indicated. **F-I,** MicroCT-based analysis of body composition in 5–6-month-old mice. **F,** Region of interest extending from proximal L1 to distal L5. **G,** Delineation of the abdominal region of interest (red), using the abdominal wall as a reference to distinguish visceral from subcutaneous adipose tissue. **H,** Plot of tissue volume across the density range used to separate adipose and lean tissues; the valley between the two peaks served as the threshold for tissue discrimination. **I,** Representative MicroCT images of male and female offspring born to dams fed either a high-fat or regular diet.

To determine whether maternal HFD exposure altered body composition in adulthood, we analyzed Off-RD and Off-HFD mice by EchoMRI (**Fig. 1E**). Adult Off-HFD mice of both sexes showed a tendency toward increased adiposity, although the difference reached statistical significance only in males (**Fig. 1E**). To further assess whether maternal HFD influences adipose tissue distribution, we performed abdominal micro-CT imaging in adult offspring. This analysis revealed increases in both visceral and subcutaneous adipose tissue (colored yellow) in male and female Off-HFD mice compared with Off-RD controls (**Fig. 1F–I**).

### Whole-body metabolic alterations in the offspring of HFD-fed mothers

We have previously reported that despite Off-HFD mice showing consistent increase in body weight and percentage of fat, their food intake and activity levels do not differ from Off-RD counterparts (21). Here, we assessed the effects of maternal obesity on whole-body metabolic parameters in adult 6-month-old Off-RD and Off-HFD mice. This assessment revealed no significant changes in the 24-hour pattern of oxygen consumption (**Fig. 2A-B**), but did illuminate significant alterations in respiratory exchange ratio (RER, **Fig. 2C-D**), an indicator of substrate utilization across active and resting periods. In Off-RD mice, the 24-hour oscillation in RER alternated between high values (utilization of carbohydrates) in active periods and low values (utilization of fats) during resting periods (**Fig. 2C-E**). These oscillations were completely absent in Off-HFD mice (**Fig. 2C-D, F**), suggesting metabolic inflexibility; correspondingly, the average 24-hour RER was significantly lower in Off-HFD vs. Off-RD (**Fig. 2G**).

**Figure 2.**
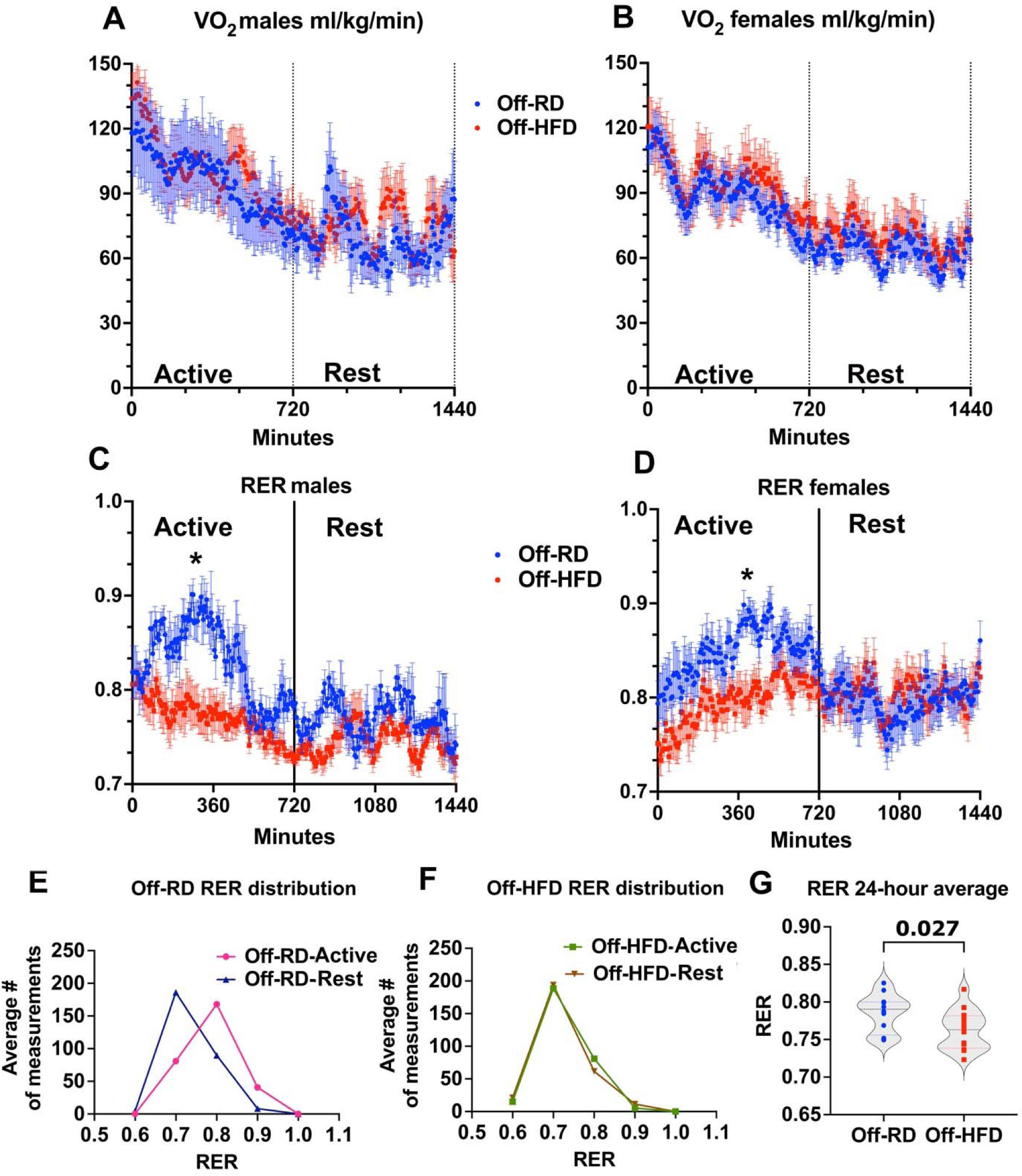
Maternal high-fat diet feeding changes metabolic phenotype in offspring. **A-G,** Oxygen consumption and respiratory exchange ratio (RER) in adult male and female Off-RD and Off-HFD mice. **A-B,** Oxygen consumption curves calculated for male (**A**) and female (**B**) mice over a 24-hour period (12 hours active and 12 hours rest). **C-D,** RER in male (**C**) and female (**D**) Off-RD and Off-HFD mice over the active and rest periods. **E-F,** Shift in RER between active and rest periods in Off-RD (**E**) and Off-HFD (**F**) mice. Male and female data are pooled. *, *p*<0.05 calculated by multiple unpaired *t*-tests with false discovery rate (FDR)<0.01. G, Average 24-hour RER in Off-RD and Off-HFD, male and female data pooled. Data are represented as individual values with lines at mean with SEM; *p*-values are shown, calculated by *t*-test. N=5–8/group of maternal diet/sex.

### Dyslipidemia and cardiac perturbations in the offspring of HFD-fed mothers

Preferential lipid utilization in offspring exposed to maternal obesity suggests metabolic reprogramming toward greater dependence on lipids as fuel. This shift is commonly associated with broader disturbances in systemic lipid handling, including enhanced lipid mobilization and impaired clearance of circulating lipids (24). To explore this association, we measured circulating levels of lipids in the blood of adult Off-RD and Off-HFD (**Fig. 3**); this revealed significantly increased levels of total cholesterol, triglycerides, phospholipids, and non-esterified fatty acids in both male and female Off-HFD mice relative to Off-RD controls (**Fig. 3A-D**). Because elevated circulating lipids are linked to altered cardiac lipid handling and adverse myocardial remodeling (25), we subsequently examined whether the dyslipidemia observed in Off-HFD offspring was accompanied by progressive cardiac dysfunction (**Fig. 4**). At three weeks of age, newly weaned Off-HFD mice exhibited a significant 8.6% increase in heart weight-to-body weight ratio compared with age-matched Off-RD mice (*p*<0.05, **Fig. 4A**). By six months of age, this difference became much more pronounced, with Off-HFD mice showing an approximately 60% increase in heart weight-to-body weight ratio, consistent with a progressive hypertrophic phenotype (**Fig. 4B**). In contrast, no differences were observed in liver weight-to-body weight ratio (**Fig. 4C**).

**Figure 3.**
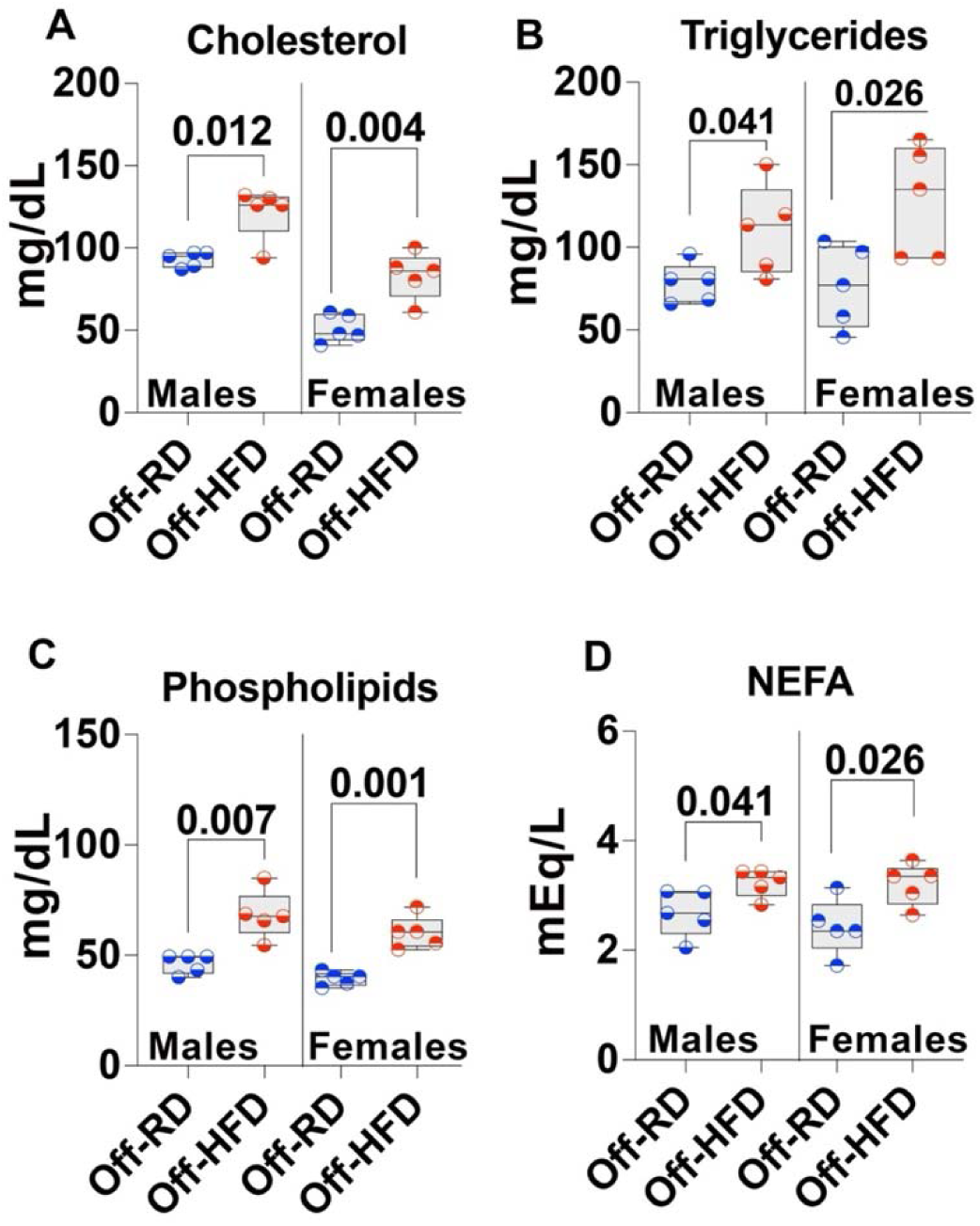
Changes in blood lipids in adult offspring of obese mothers. **A,** Total cholesterol. **B,** Triglycerides. **C,** Phospholipids, and **D,** Non-esterified fatty acids (NEFA). Data were analyzed by multiple *t*-tests with FDR<0.01 and are represented as box and whiskers with minimum and maximum and individual values with lines at mean. N=5/sex/group of maternal diet, 20 animals overall; *p*-values are shown.

**Figure 4.**
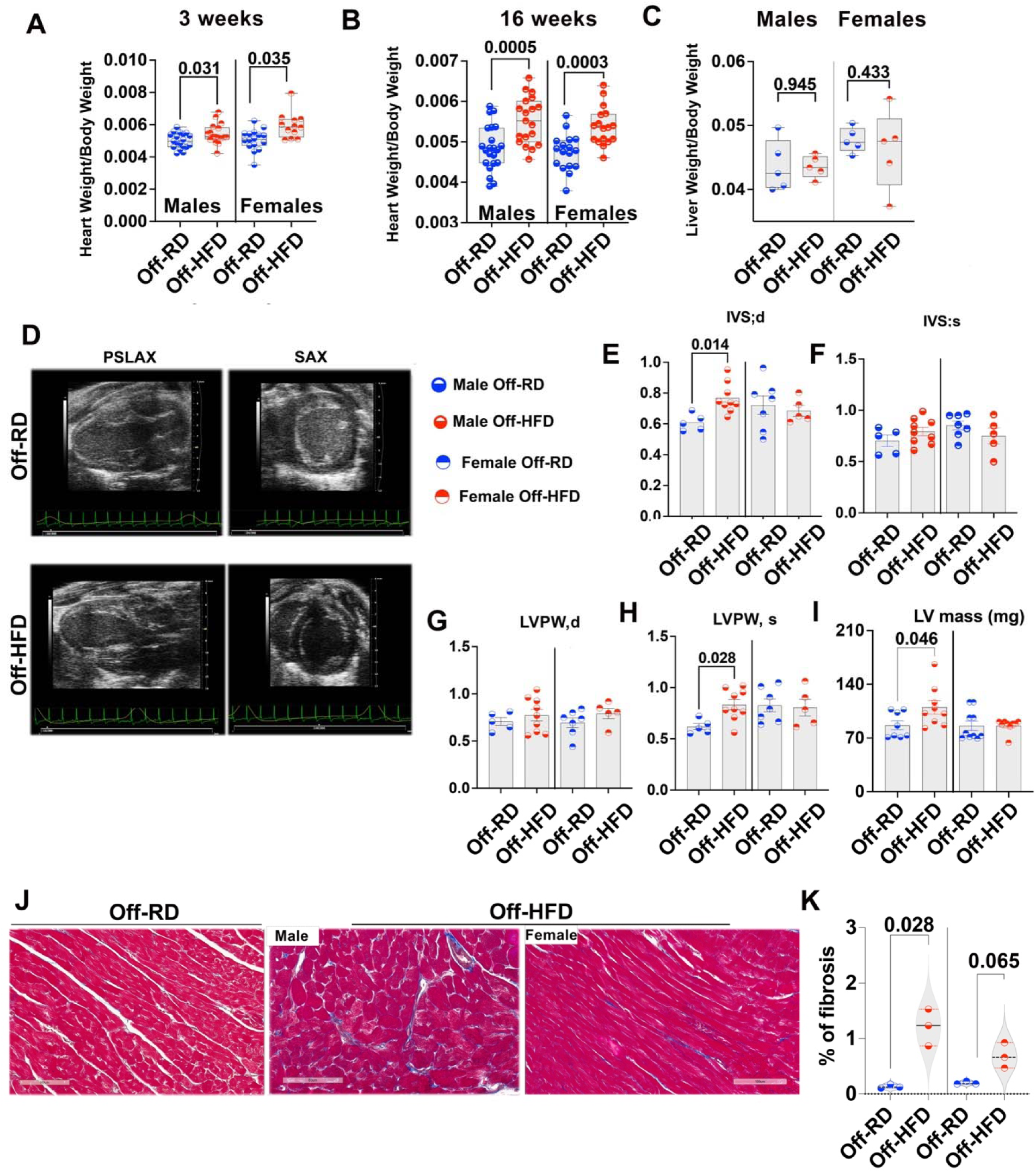
Cardiac hypertrophy in the offspring of obese mothers. **A-B,** Heart weight-to-body weight ratio in newly weaned (**A**) and six-month-old (**B**) Off-RD and Off-HFD mice fed a normal chow diet. N=38–62 mice/group of maternal diet. **C,** Liver weight-to-body weight ratio, N=10/experimental group. **D,** Representative parasternal longLaxis BLmode images of the left ventricle and short axis. **E-I,** Sex-dependent changes in diastolic (**E**) and systolic (**F**) intra-ventricular septum thickness; and in diastolic (**G**) and systolic (**H**) left ventricular posterior wall thickness and LV mass (**I**) in male and female Off-RD and Off-HFD. N=5–9/sex/group of maternal diet. Data are represented as individual values for males (left) and females (right) with lines at mean with SEM. Significant differences were determined by Mann-Whitney test followed by unpaired *t*Ltest; *p*-values are shown. **J,** Masson’s trichrome staining of cardiac sections. Scale bars: 100 μm (Off-RD) and 100 and 80 μm (Off-HFD). **K,** Quantification of fibrosis. Blue-stained areas in trichrome staining were color-thresholded using NIH ImageJ and quantitated. Data are represented as individual values with lines at mean with SEM. N=3/sex/group of maternal diet; *p*-values are shown.

To further assess cardiac structure and function, we performed echocardiography in six-month-old male and female Off-HFD and Off-RD mice (**Fig. 4D-I**). Although most echocardiographic parameters were not significantly different between groups, these analyses revealed a sexually dimorphic cardiac response to maternal obesity. Specifically, male but not female Off-HFD mice showed increased interventricular septal thickness in diastole (IVS;d, **Fig. 4E**), left ventricular posterior wall thickness in systole (LVPW;s, **Fig. 4H)**, and LV mass (**Fig. 4I**), consistent with left ventricular hypertrophy.

Pathological cardiac hypertrophy is often accompanied by fibroblast activation and extracellular matrix accumulation, including collagen deposition (26). Therefore, we then determined whether the hypertrophic Off-HFD hearts also exhibited evidence of cardiac fibrosis by performing histological analysis of trichrome-stained paraffin sections from adult Off-RD and Off-HFD hearts (**Fig. 4J-K**). This revealed a 2.5-fold increase in interstitial fibrosis in male Off-HFD hearts compared with Off-RD controls. Female Off-HFD hearts showed a trend toward increased fibrosis that did not reach statistical significance (*p*=0.065, **Fig. 4K**).

From these findings, we hypothesized that progressive cardiac hypertrophy in Off-HFD offspring may reflect concurrent vascular remodeling (27). To address this, we studied whether maternal obesity also alters arterial stiffness in the offspring. We therefore measured pulse wave velocity (PWV) in male and female Off-RD and Off-HFD mice (**Fig. 5**). PWV was significantly increased in six-month-old male Off-HFD mice compared with age-matched Off-RD controls (*p*<0.05, **Fig. 5A-B**), whereas no changes in vascular stiffness were observed in females. These findings indicate that cardiac remodeling in male Off-HFD offspring is accompanied by vascular dysfunction. Increased arterial stiffness has been shown to elevate central systolic pressure and left ventricular afterload, thereby promoting compensatory hypertrophic growth (28); moreover, reduced arterial compliance is a well-established contributor to the development of hypertension (29). Consistent with this interpretation, measurement of mean arterial pressure by tail-cuff sphygmomanometry revealed a significant increase in adult six-month-old male Off-HFD mice, while no change was detected in female Off-HFD mice (**Fig. 5C**).

**Figure 5.**
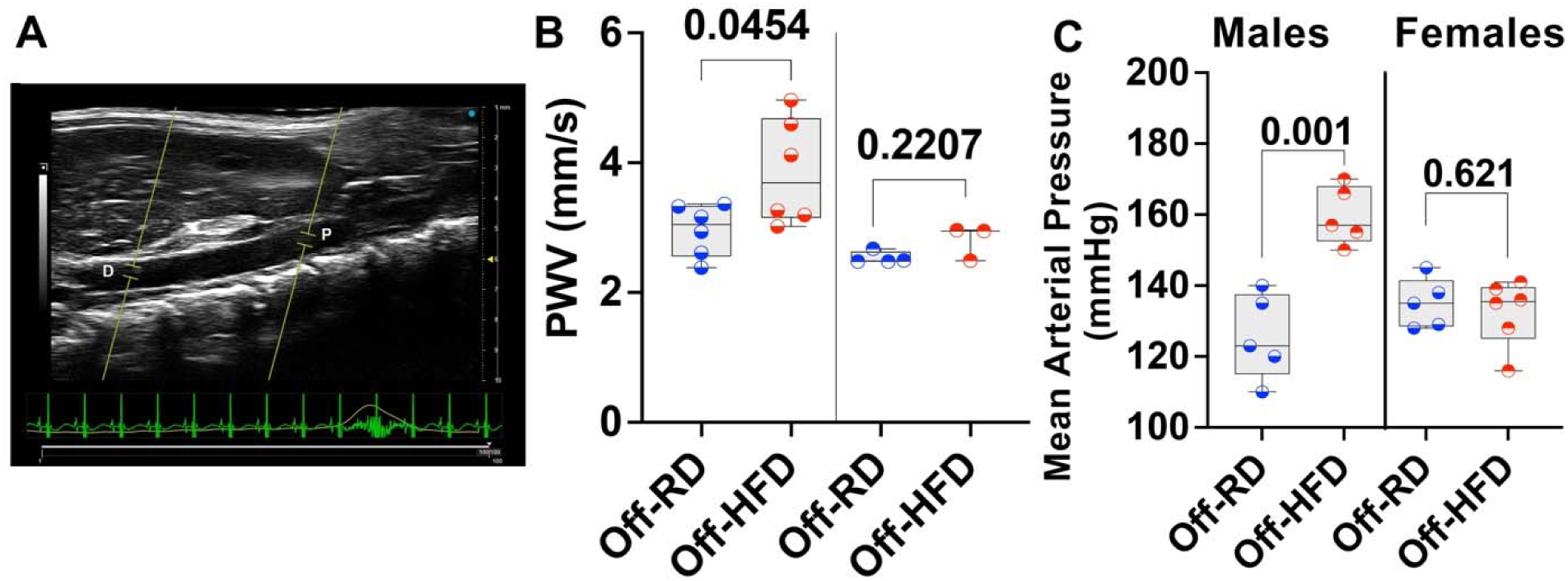
Maternal obesity increases arterial stiffness in male offspring. **A-B,** Representative ultrasound image (**A**) and calculations of pulse wave velocity (PWV) (**B**) in 6-month-old male and female Off-RD and Off-HFD. Blood flow waveforms were continuously recorded for 20 sec. at a proximal and a distal location along the abdominal aorta with simultaneous ECG. PWV was calculated by dividing the distance between the proximal and distal locations by the difference between the proximal (P) and distal (D) transit times and expressed in mm/sec. N=3–6/sex/experimental group. **C,** Tail cuff-measured mean arterial pressure in 6-month-old male and female Off-RD and Off-HFD mice. Data are represented as box and whiskers with minimum and maximum and individual values with lines at mean. Statistical analysis was performed using two-way ANOVA followed by *t*-test. N=5–6 mice/sex/group of maternal diet; *p*-values are shown.

### Ultrastructural and metabolic alterations in the hearts of adult Off-HFD mice

Cardiac hypertrophy in adult Off-HFD offspring was accompanied by marked ultrastructural abnormalities, as revealed by transmission electron microscopy (**Fig. 6A**). In control hearts, mitochondria were densely packed in an organized array between sarcomeres, with their transverse alignment closely matching the Z-lines. In contrast, Off-HFD hearts displayed disrupted mitochondrial architecture, with mitochondrial bundles that lacked this regular spatial organization. Off-HFD hearts also showed an accumulation of autophagic structures, including autophagosomes and autolysosomes (white asterisks), together with increased number of lipid droplets (yellow arrows). To determine whether these structural abnormalities associated with impaired mitochondrial function, we assessed the expression of representative protein subunits from all five electron transport chain complexes by western blotting (**Fig. 6B-H**). Although no sex-specific differences were detected (**Figure S1**), pooling of data from males and females demonstrated significant reductions in expression of complexes I, III, and V in Off-HFD hearts. Complex IV also showed a downward trend, although this did not reach statistical significance (*p*=0.066), whereas complex II remained unchanged.

**Figure 6.**
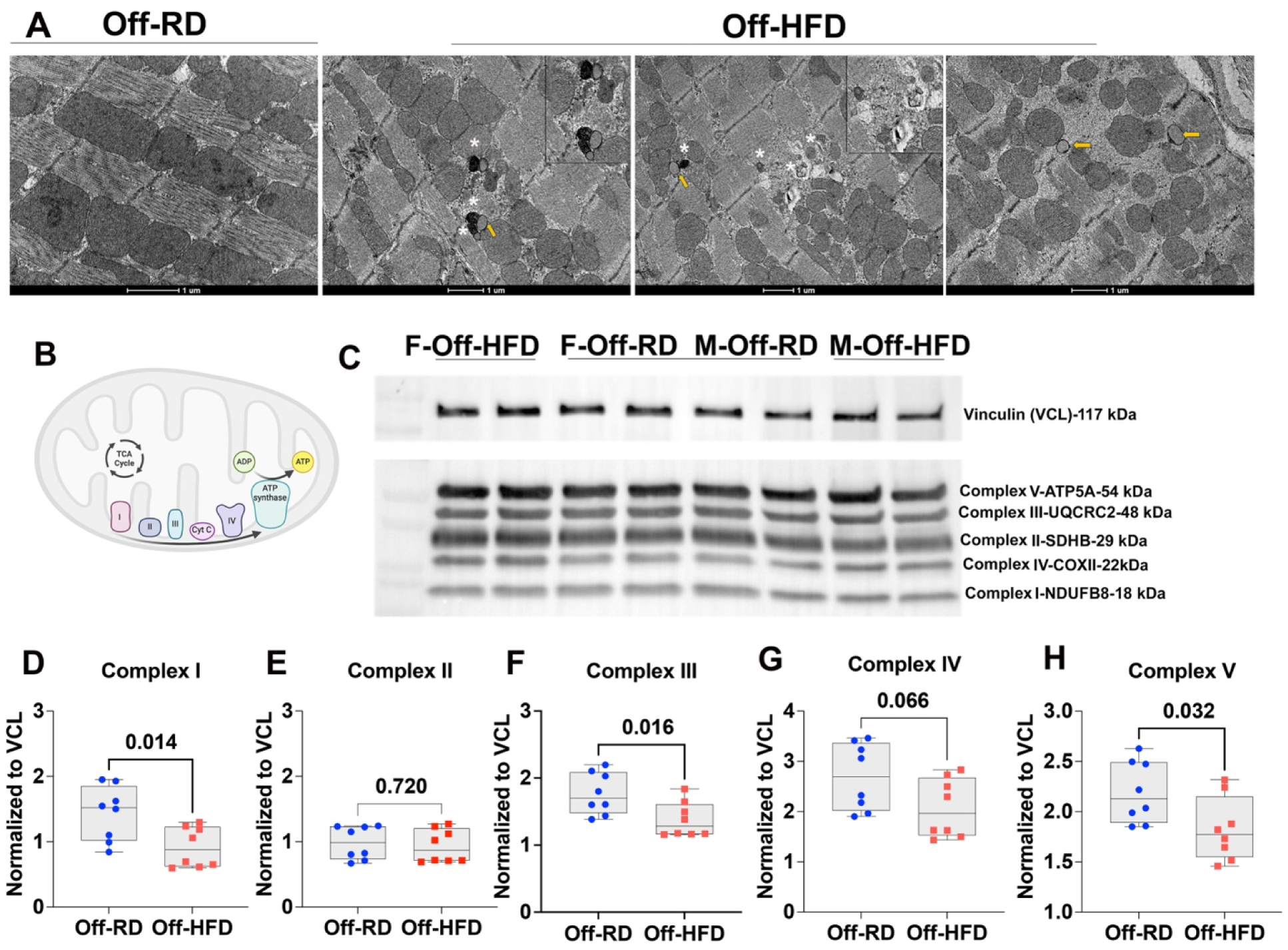
Ultrastructural and metabolic changes in Off-HFD hearts. **A,** Transmission electron microscopy from adult mice showing dysregulations in mitochondrial alignment and increased accumulation of autolysosomes (asterisks) and lipid droplets (yellow arrows) in Off-HFD vs. Off-RD. Inserts present magnified views of autophagic vacuoles and lipid droplets. Scale bar: 1 μm. **B-H,** Western blots for expression of subunits of the mitochondrial electron transport chain, with Vinculin as loading control. **B,** Cartoon of the mitochondrial complexes, created with Biorender.com. **C,** Representative western blots for male and female Off-RD and Off-HFD. **D-H,** Quantification data for complex I (**D**), II (**E**), III (**F**), IV (**G**), and V (**H**). Data were analyzed by two-way ANOVA followed by *t*-test and are represented as box and whiskers with minimum and maximum and individual values with lines at mean. N=8/group of maternal diet; *p*-values are shown. Data from males and females were pooled.

These ultrastructural findings were supported by our *in vivo* label-free photoacoustic imaging (PAI, **Fig. 7A**), which allowed us to visualize accumulation of lipids, average oxygen saturation (sO_2_Av) and total hemoglobin. As a point of comparison, we additionally imaged hearts from six -month-old mice that were fed the HFD for eight weeks to develop diet-induced obesity (**Fig. 7B**). This comparison revealed that, despite being fed the RD only, male and female Off-HFD mice presented an increase in cardiac lipid accumulation along with reduced oxygenated hemoglobin signal in a sex-independent manner similar to the HFD-fed obese animals. Among the offspring groups, lipids were increased by 45% from 0.067 in Off-RD to 0.147 in Off-HFD (arbitrary units), and sO_2_Av was decreased markedly from 72.15% in Off-RD to 63.25% in Off-HFD (*p*<0.05). No change was observed in total hemoglobin (**Fig. 7C-E**). Overall, the concordance of these imaging and ultrastructural observations suggests that maternal obesity promotes ectopic and intracellular lipid deposition within the heart, which may contribute to mitochondrial dysfunction and metabolic stress.

**Figure 7.**
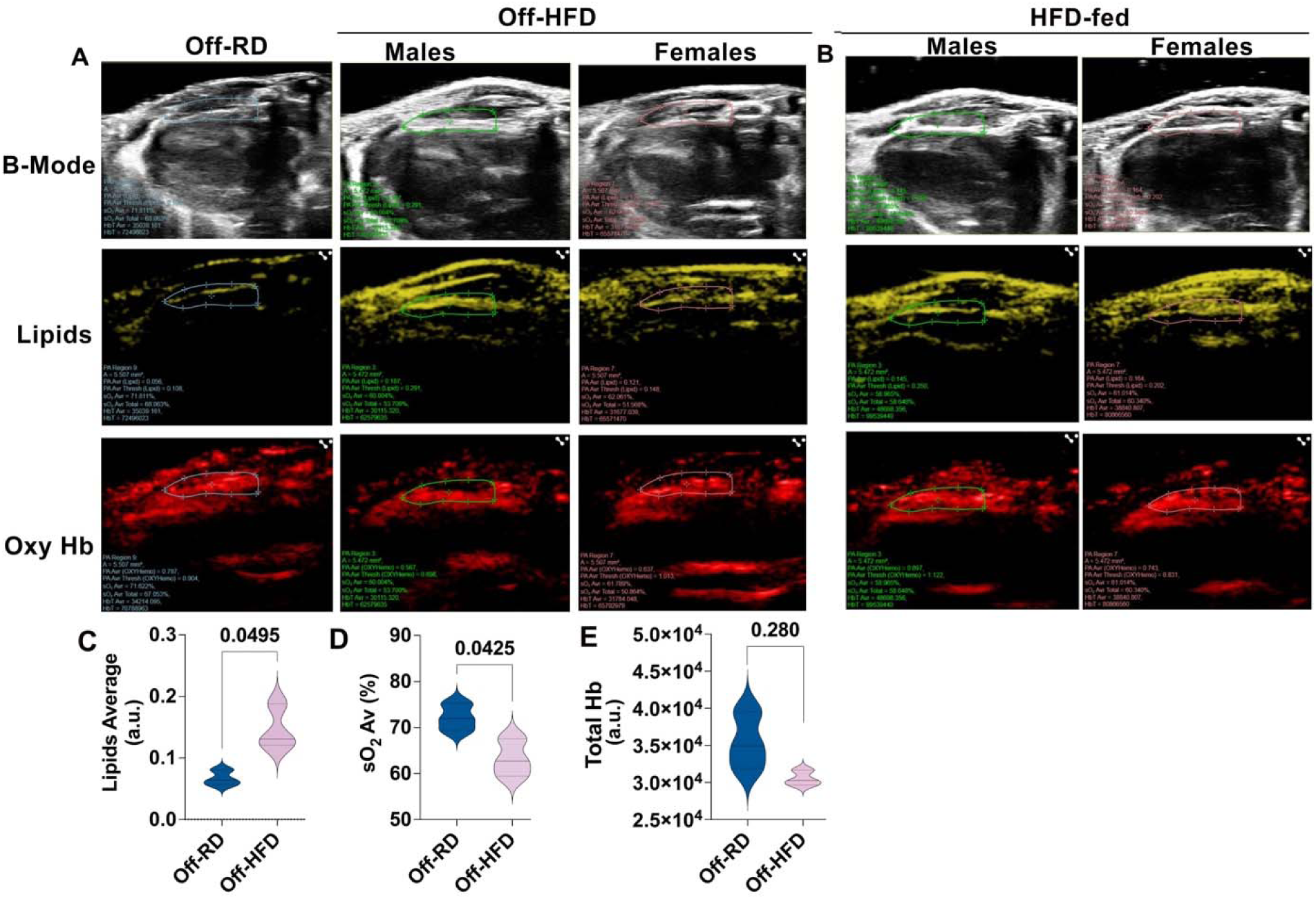
Photoacoustic assessment of Off-RD and Off-HFD hearts at 4 months of age. **A,** Ultrasound-guided photoacoustic images displaying B-mode (top panel) overlayed with lipid accumulation (middle panel) and oxygen saturation (lower panel) in male and female offspring of RD and HFD-fed dams. **B,** Corresponding assessments done in male and female mice fed the regular or high-fat diet for 8 weeks. **C-E,** Quantitative data presented as volume of lipid-rich areas (**B**) and total hemoglobin (both in arbitrary units, D) and oxygenation levels in % (**C**). Wavelengths: 1210 nm for lipids, 750 nm for oxyhemoglobin, 850 for deoxyhemoglobin (total hemoglobin). Data from males and females were pooled. N=3-4/sex/experimental group; *p*-values are shown. Scale bar: 2 mm.

Since mitochondrial abnormalities are closely linked to activation of autophagy (30), and because we observed ultrastructural evidence of autophagic vacuole accumulation in Off-HFD hearts, we further assayed marker proteins respectively involved in the early (TFEB), intermediate (p62), and late (Rubicon) stages of autophagic progression, along with the autophagy-related markers ATG7 and LC3-II (cleaved LC3). Unexpectedly, no significant between-group differences were observed. However, ATG7 expression was higher in female than in male hearts, independent of maternal diet (**Figure S2**).

### Immune perturbations in adult Off-RD and Off-HFD hearts

Since mitochondrial dysfunction is a potent driver of cardiac inflammation (31), we next asked whether maternal obesity alters the immune landscape of the offspring heart. To address this question, we performed flow cytometric profiling of cardiac immune populations in adult male and female Off-RD and Off-HFD mice; the gating strategy is depicted in **Figure S3**. Female Off-HFD hearts displayed a reduction in total CD45+ leukocytes compared with female Off-RD controls, whereas total CD11b+ cells were unchanged across groups (**Fig. 8A-B**). Notably, despite the stable overall CD11b+ population, female Off-HFD hearts exhibited significant increases in both CD11b+F4/80+ monocyte-derived macrophages and CD11b−F4/80+ resident macrophages; no comparable changes occurred in males (**Fig. 8C-D**). Total CD3+ T cells tended to be reduced in both male and female Off-HFD hearts, although these differences did not reach statistical significance (**Fig. 8E**). In females, this was accompanied by significant reductions in both CD4+ and CD8+ T-cell subsets relative to Off-RD controls (*p*<0.05, **Fig. 8F-G**). Meanwhile, double-negative T-cell abundance was not altered by maternal diet in either sex (**Fig. 8H**).

**Figure 8.**
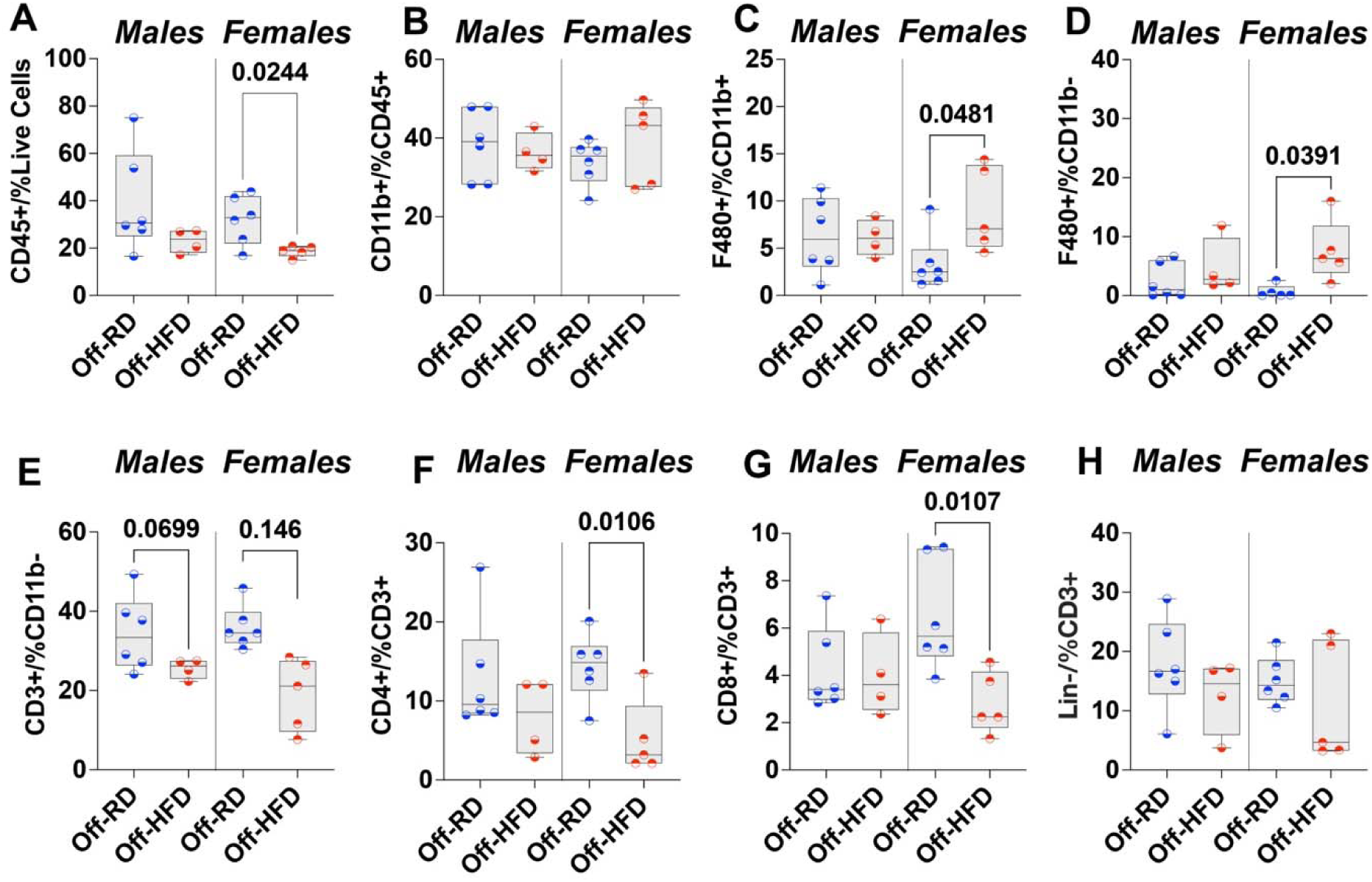
Flow cytometry for immune cell populations in hearts of adult male and female Off-RD and Off-HFD. **A,** CD45+; **B,** CD11b+; **C,** F480/CD11b+; **D,** F480/CD11b-; **E,** CD3+ T-cells; **F,** CD4+T cells; **G,** CD8+ T-cell; **H,** double negative T-cells. Data are represented as box and whiskers with minimum and maximum and individual values with lines at mean N=4–6/sex/group of maternal diet; *p*-values are shown.

Taken together, our findings demonstrate that maternal obesity programs a progressive cardiac phenotype in offspring that is characterized by early-life hypertrophic growth and subsequent lipid accumulation, fibrosis, mitochondrial dysfunction, and selective immune remodeling in adulthood. These observations raise the possibility that the transition from early cardiac growth changes to overt adult pathology is partly driven by persistent epigenetic reprogramming established during development. We next investigated whether cardiac phenotype is associated with temporal changes in cardiac DNA methylation by performing DNA methylation analysis in hearts from three-week-old and six-month-old offspring. The early hypertrophic phenotype observed at three weeks may reflect initial epigenetic programming events, whereas the more severe hypertrophy and associated metabolic, structural, and immune abnormalities observed at six months may be linked to persistence or amplification of these changes. In parallel, we performed RNA sequencing on the same six-month-old hearts to identify transcriptional pathways associated with the adult disease phenotype. This approach allowed us to define the epigenetic and transcriptional trajectory underlying maternal obesity-induced cardiac remodeling.

### DNA methylation changes in the left ventricle of newly weaned Off-HFD mice

To identify regions and individual CpGs with differential DNA methylation between offspring of HFD- and RD-fed mothers, we performed RRBS on left ventricular tissue collected from newly weaned, three-week-old male and female Off-RD and Off-HFD mice. To assess sample similarity based on global methylation profiles, we conducted principal component analysis using methylation percentages from 343,296 CpGs with at least 10X coverage; the corresponding plot is shown in **Figure S4**. One male Off-HFD sample was excluded because of low coverage. In the initial analysis, three samples appeared to segregate away from the cluster formed by the rest; these few also showed the highest methylation in non-CpG contexts. When those segregated samples were excluded, the plot showed female Off-RD to cluster with female Off-HFD, and males to similarly cluster together (**Figure S4)**. Next, we identified DMRs with an absolute methylation difference greater than 10% and a Šídák-adjusted *p*-value <0.05. Using these criteria, we detected 61 DMRs located within protein-coding genes (**Fig. 9A**), including 13 hypermethylated and 48 hypomethylated regions in Off-HFD hearts. The top ten hypermethylated and hypomethylated DMRs, along with their associated genes, are listed in **Tables S1** and **S2**. To assess the biological relevance of these methylation changes, we performed STRING protein–protein interaction network analysis on the genes associated with DMRs (**Fig. 9B**). *Notch1,* a key regulator of cardiac development that also contributes to cardiac hypertrophy and fibrosis (32), emerged as a central interacting gene within this network. More specifically, *Notch1* connected with several genes relevant to cardiac structure, metabolism, fibrosis, and inflammation, including *Epas1,* which encodes HIF-2α and is essential for normal mitochondrial function in the heart (33); Endoglin (*Eng*), a regulator of cardiac fibrosis (34); Platelet endothelial cell adhesion molecule-1 (*Pecam1/CD31*), which mediates inflammatory responses (35); and CCAAT/enhancer-binding protein alpha (*Cebpa*), which is activated in the heart in response to both developmental cues and injury signals (36). We further performed Gene Ontology (GO) term enrichment analysis using EnrichR to identify pathways associated with differentially methylated genes (**Fig. 9C**). This revealed significant enrichment of developmental cardiac pathways, including endocardial cushion formation, cardiac chamber morphogenesis, cardiac septum morphogenesis, and heart trabeculation.

**Figure 9.**
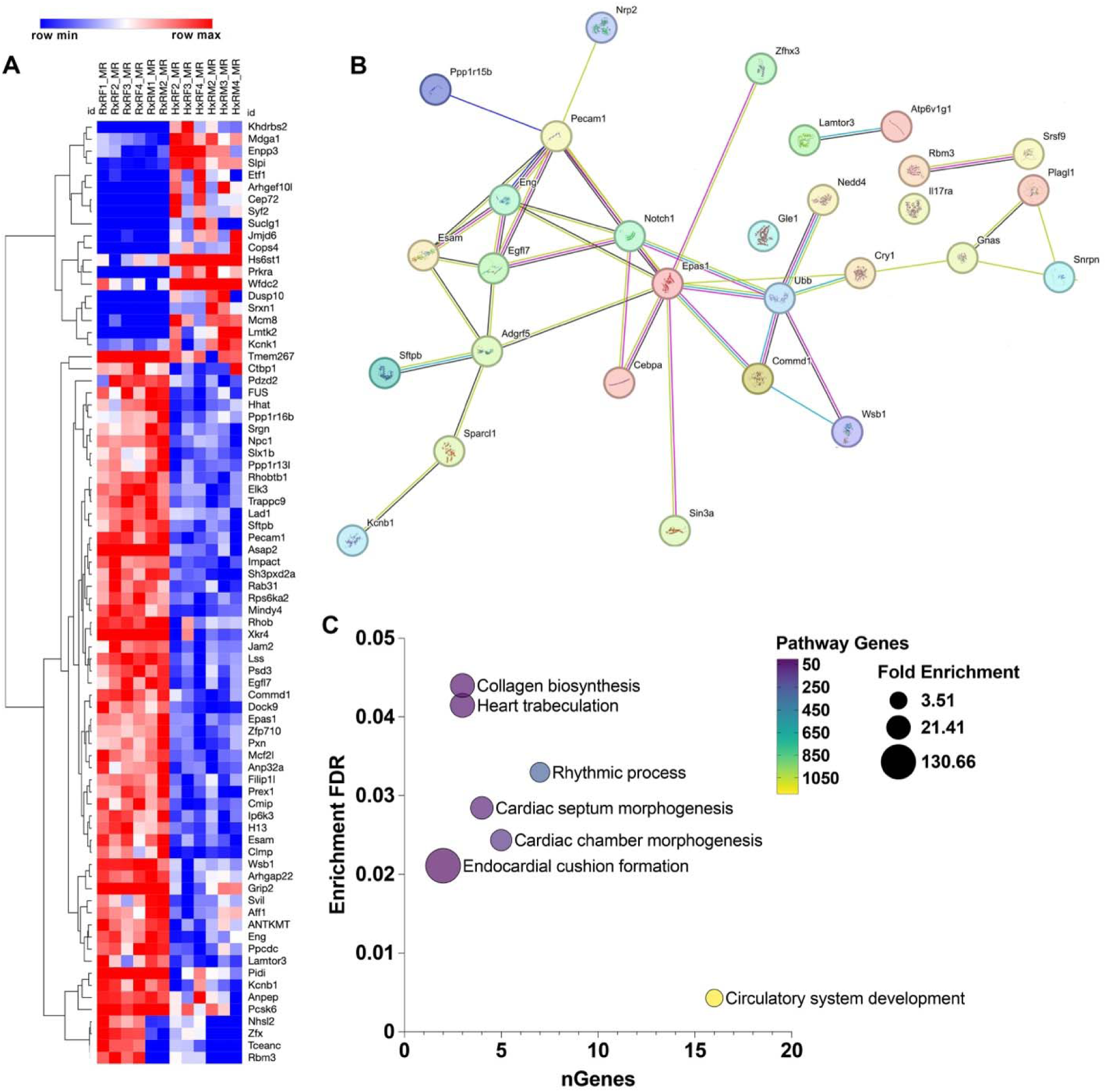
Mapping and comparisons of the DNA methylomes of newly weaned Off-RD and Off-HFD. **A,** Heatmap and unsupervised hierarchical clustering of male and female Off-RD and Off-HFD samples based on individual DNA methylation status. **B,** Mapping of differentially methylated genes using STRING. Colored nodes: genes with differentially methylated CpGs and first shell of interactors. **C,** Bubble chart showing Gene Ontology biological processes enriched among genes with differentially methylated regions.

### DNA methylation changes in adult Off-HFD mice

To determine whether epigenetic alterations induced by maternal obesity early in life persist into adulthood, we next performed DNA methylation analysis on left ventricles from six-month-old Off-RD and Off-HFD mice (**Fig. 10**). We constructed multidimensional scaling plots to visualize similarities between samples in an unsupervised manner (**Figure S5**), which showed distinct separation by sex and less-distinct separation by maternal diet group with no major outliers. Manhattan plots of CpGs associated with maternal obesity in males and females are illustrated in **Figure S6A–B**. CpG islands were found to be preferentially hypomethylated in male Off-HFD mice and hypermethylated in female Off-HFD mice (**Figure S6C-D**). Conversely, CpG shores and open sea regions were predominantly hypermethylated in male Off-HFD mice and hypomethylated in female Off-HFD mice.

**Figure 10.**
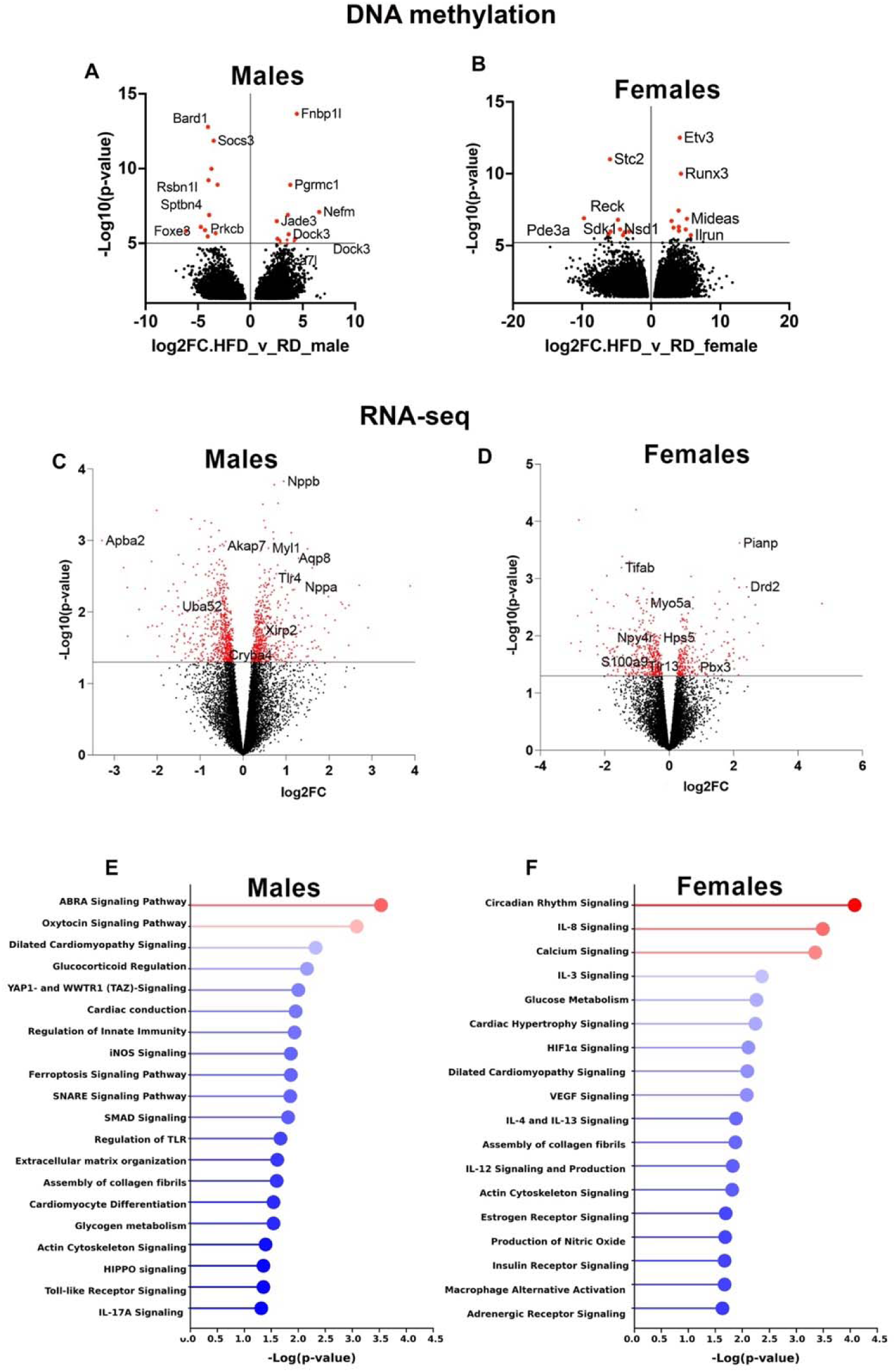
DNA methylation and RNA-seq data in the hearts of 6-month-old male and female Off-RD and Off-HFD. **A-B,** Volcano plots for DNA methylation data for male (**A**) and female (**B**) Off-HFD showing statistical significance (-Log(*p*-value)) vs. magnitude of change (Log2FC). Dotted lines indicate the threshold for significance. **C-D,** Mapping and comparisons of RNA-seq data from the left ventricles of the same animals as in A-B. Volcano plots for DEGs in male (**C**) and female (**D**) Off-HFD showing statistical significance (-Log(*p*-value)) vs. magnitude of change (Log2FC). Dotted lines indicate the threshold for significance. **E-F,** Canonical pathways identified by Ingenuity Pathway Analysis as enriched in left ventricles of male (**C**) and female (**D**) Off-HFD mice. N=5/sex/group of maternal diet.

Using a cutoff of *p* < 0.05 and absolute fold change >1.5, we identified a respective 20,585 and 20,422 hypomethylated CpG sites in male and female Off-HFD mice, along with 26,257 and 27,607 hypermethylated sites (**Fig. 10A–B**). In both males and females, promoter regions were preferentially represented among differentially methylated sites compared with distal genomic regions, including exons, introns, and 5′-UTRs (**Fig. S5E–F**).

Subsequently, we performed gene set enrichment analysis using the genes nearest to each differentially methylated CpG. Regions exhibiting consistent differential methylation in Off-HFD mice at both three weeks and six months of age were associated with 19 genes (**Fig. S7**). Of these, 11 have previously been linked to cardiovascular disease, and five have been associated with lipid metabolism (**Table S3**). The direction of DNA methylation changes was preserved for 10 genes across mice aged three weeks and six months. Eight genes, Cmip,Eng, Esam, Mindy4, Pcsk6, Sh3pxd2a, Tmem267, Rab31, shifted from hypermethylation at three weeks to hypomethylation at six months, and one gene, Stag2, exhibited the opposite pattern.

### Transcriptomic changes in adult offspring

To determine whether persistent DNA methylation changes were accompanied by altered gene expression, we performed RNA sequencing on the same hearts used for methylation analysis in adult mice. Volcano plots showing up- and downregulated differentially expressed genes (DEGs) in male and female Off-HFD hearts are presented in **Fig. 10C–D**. Maternal obesity induced a marked sex-dependent transcriptomic response. In male Off-HFD hearts, 176 genes were upregulated and 197 were downregulated relative to male Off-RD hearts (*p* < 0.05, |FC| > 1.5). In female Off-HFD hearts, 106 genes were upregulated and 200 were downregulated using the same criteria. Only 52 DEGs were shared between males and females; notably, all shared genes were regulated in opposite directions between the sexes (**Figure S8**).

A total of 96 DEGs in male and female Off-HFD hearts also showed corresponding changes in DNA methylation regions (**Table S4**). Several of those genes were related to the cardiac conduction system (*Kcnh2*, *Kcnd3*, and *Scn4b*). Other identified genes were associated with lipid metabolism (*Dgkb*, *Sgms1*, *Stard3*), mitochondrial physiology (*Mfn2*, *Mrpl37*, *Mrps35*), nuclear organization (*Lamb1*, *Lmnb1*), and epigenetic regulation (*H4c9*, *Hdac9*).

To determine whether transcriptional changes in Off-HFD hearts were enriched in specific biological pathways, we utilized Ingenuity Pathway Analysis (IPA) (**Fig. 10E–F**). In male Off-HFD hearts, differentially regulated pathways included actin-binding rho activating protein (ABRA) signaling, a biomechanical stress sensor (37), pathways associated with extracellular matrix organization, and YAP/TAZ and Hippo signaling (**Fig. 10E**). Male Off-HFD hearts also showed enrichment of immune-related pathways, including IL-17A signaling, toll-like receptor signaling, and regulation of innate immunity.

In female Off-HFD hearts, significantly enriched pathways included those related to circadian rhythm, cardiac hypertrophy and dilated cardiomyopathy, HIF-1 signaling, and regulation of the actin cytoskeleton (**Fig. 10F**). Similar to males, female Off-HFD hearts also showed enrichment of inflammation-related pathways, including macrophage alternative activation and interleukin signaling pathways involving IL-3, IL-4, IL-8, IL-12, and IL-13.

IPA upstream regulator analysis further revealed sex-dependent enrichment of miRNAs and transcriptional regulators among DEGs (**Table S5**). In male Off-HFD hearts, two transcription factors—*Smad3*, a regulator of cardiac fibrosis (38), and *Glis3*, a gene involved in embryonic development (39)—exhibited an approximately five-fold increase in expression. Congruently, the corresponding signaling pathways were predicted to be activated in male Off-HFD hearts (z-score > 2). *Ppargc1b*, a key transcriptional regulator of mitochondrial function and lipid utilization, was also predicted to be significantly activated (z-score > 2). Of regulators predicted to be inhibited, nine miRNAs with known roles in cardiac hypertrophy were identified in male Off-HFD hearts: miR-1-3p (40), miR-27a-3p (41), mir-8 (42), miR-29b-3p (43), miR-30c-5p (44), mir-140 (45), miR-124-3p (46), mir-9 (47), and miR-34a-5p (48).

In female Off-HFD hearts, expression of the transcription factor *Meis1*, a regulator of the cardiac conduction system, was increased 1.5-fold vs. female Off-RD hearts. In addition, ARNT and FOXO3 signaling, associated with cardiac lipid metabolism (49) and autophagy (50) respectively, were predicted to be significantly activated. Among regulators predicted to be inhibited, we observed three miRNA pathways—miR-210, miR-30, and miR-135—associated with cardiac mitochondrial function (51), cardiac hypertrophy (52), and fibrosis (53), respectively. In addition, three transcription factors (*Etv4*, *Lmo2*, and *Pax6*) exhibited significantly altered expression (FC=-6.31, 3.08, 1.80), while their corresponding pathways were inhibited. In keeping with the pathway analysis, BMAL1 signaling, a regulator of circadian rhythm (54), was reduced in female Off-HFD hearts. Together, the RNA-seq data demonstrate that maternal obesity induces extensive and sex-dependent transcriptional changes in the hearts of adult offspring.

To identify potential upstream regulatory mechanisms underlying these changes, we examined DNA methylation from six-month-old Off-HFD hearts using Enrichr-based transcription factor protein–protein interaction (PPI) analysis (**Figure S9**). In male Off-HFD hearts, the PPI analysis identified several potentially important factors, including estrogen receptor alpha (ESR1), a nuclear hormone receptor increasingly recognized as an important regulator of cardiac remodeling, hypertrophic gene expression, myocardial metabolism, fibrosis, angiogenesis, and sex-dependent responses to cardiac stress (55, 56). In both male and female Off-HFD hearts, c-Myc (MYC) was affected; this transcription factor is involved in developmental and growth-related programs and known to be induced in the heart by stress signals, including hypertrophic stimuli (57).

We also utilized IPA to integrate the six-month RRBS and RNA-seq datasets and identify shared regulatory submodules that may contribute to the cardiac phenotype observed in Off-HFD offspring (**Table S6**). Notably, in male Off-HFD hearts, the predicted activated pathways were predominantly pro-inflammatory and included TNFα, TLR4, NF-κB, and IL-17A, whereas ADAM12 and miR-30c-5p pathways, previously associated with cardioprotection (58, 59), were predicted to be inhibited. Conversely, female Off-HFD hearts showed predicted inhibition of several pro-inflammatory regulators, including IFNG, IL-5, IL-4, and TNF, together with activation of Nuclear factor of activated T cells 5 (*Nfat5*), a stress-responsive transcription factor involved in cardiovascular development and calcium signaling, and also recently linked to suppression of cardiac fibrosis (60); also inhibited were Protein kinase AMP-activated catalytic subunits alpha 1 and 2 (*Pprkaa1/Prkaa2*), which are responsible for driving glucose uptake and fatty acid oxidation and hence play crucial roles in cardioprotection (61). We further asked what cardiac genes in Off-HFD were differentially methylated early in life and also exhibited differential expression in adulthood (**Table S7**). Overall, 23 genes were consistently altered, including several related to lipid metabolism and obesity (*Ptpra, St3gal1, Ncoa1, Sparcl1, Copg1, Maz, Stard3*).

Since Off-HFD hearts showed evidence of cardiac hypertrophy and fibrosis, we next examined the expression of genes associated with hypertrophic growth and extracellular matrix remodeling in the interest of determining whether the broader transcriptomic alterations induced by maternal obesity included molecular signatures consistent with progression toward pathological cardiac remodeling. The results are shown in **Fig. 11**. Although both male and female Off-HFD mice exhibited cardiac hypertrophy, expression of hypertrophy-related genes was markedly influenced by offspring sex. Specifically, male Off-HFD hearts showed significant upregulation of B-type natriuretic peptide (*Nppb*) and atrial natriuretic peptide (*Nppa*), indicating reactivation of two fetal genes that are well-established markers of cardiac stress and pathological hypertrophy (62) (**Fig. 11A–B**). Male Off-HFD hearts also showed altered expression of WW domain-containing transcription regulator protein 1 (*Wwtr1/Taz*), a transcriptional co-activator in the Hippo signaling pathway (63) (**Fig. 11C**), along with other core Hippo pathway components, including Mps1 binder-related kinase activator-like 1A (*Mob1a*) (64) and TEA domain transcription factor 2 (*Tead2*), a transcriptional regulator of downstream genes involved in cell proliferation, survival, and differentiation (65) (**Fig. 11D–F**). In addition, *Tbx5*, a transcription factor whose deficiency and overexpression have both been implicated in cardiac remodeling (66), was significantly reduced in male Off-HFD hearts compared with male Off-RD hearts.

**Figure 11.**
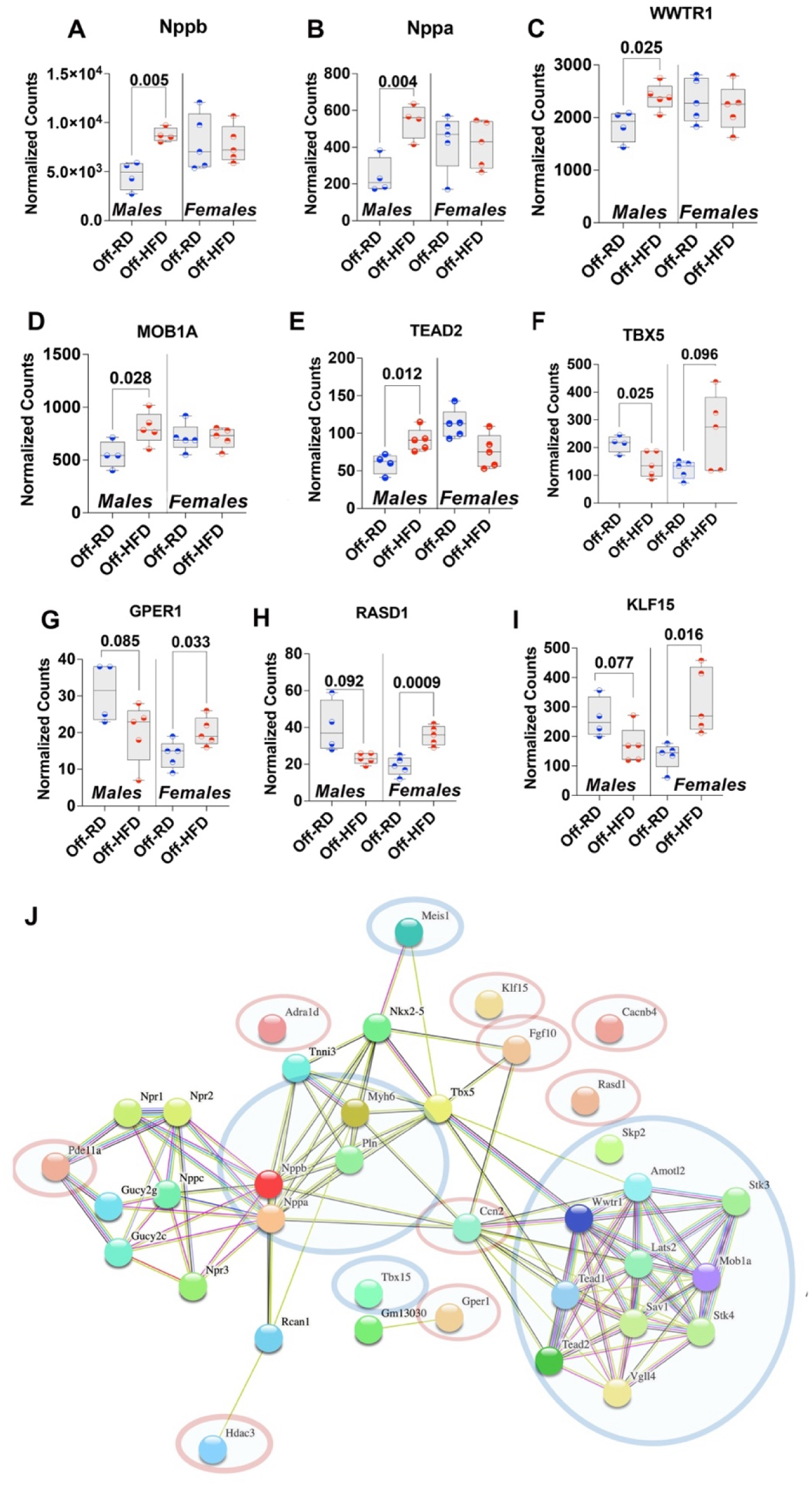
Sexual dimorphism in expression of genes associated with cardiovascular disease in adult male and female Off-RD and Off-HFD. **A-F,** Normalized counts for DEGs in male Off-HFD vs. Off-RD showing markers of heart failure *Nppb* (**A**) and *Nppa* (**B**) and members of Hippo (Yap/Taz) signaling including *Wwtr1* (**C**), *Mob1a* (**D**), *Tead2* (**E**) and *Tbx5* (**F**). **G-I**, Normalized counts for DEGs in female Off-HFD vs. Off-RD including *Gper1* (**G**), *Rasd1* (**H**), and *Klf15* (**I**). **J**, Protein interaction map of DEGs using STRING in male (blue circle) and female (red circle) Off-HFD vs. Off-RD. N=4–5/sex/group of maternal diet; *p*-values are shown.

In contrast to males, female Off-HFD hearts showed increased expression of G protein-coupled estrogen receptor 1 (*Gper1*), which has a reported cardioprotective role (67) (**Fig. 11G**). Also increased was Ras dexamethasone-induced protein 1 (*Rasd1*), whose downregulation has been linked to impaired cardiac autophagy and the progression of hypertrophic cardiomyopathy (68). This same gene was significantly downregulated in male Off-HFD hearts, suggesting that preservation or upregulation of RASD1 may contribute to a more protective cardiac response in females (**Fig. 11H**). Similarly, Krüppel-like factor 15 (*Klf15*), a regulator of cardiac lipid metabolism (69) that also acts as a negative modulator of pathological cardiac hypertrophy, fibrosis, and heart failure (70), was significantly increased in female Off-HFD hearts, while showing a trend toward decreased expression in males (**Fig. 11I**). Together, these findings indicate that maternal obesity induces sex-dependent changes in cardiac hypertrophy–associated genes, with male Off-HFD hearts showing greater activation of fetal and hypertrophic remodeling markers whereas female Off-HFD hearts exhibit increased expression of genes associated with cardioprotection and suppression of pathological hypertrophy.

To further characterize the possible functional relationships among hypertrophy-associated DEGs, we performed protein–protein interaction network analysis using the STRING database. DEGs identified in male and female Off-HFD hearts relative to their respective Off-RD controls, including genes shown in **Fig. 11A-I** as well as additional DEGs not shown, were mapped together (**Fig.11J**). Male-associated genes are indicated by blue circles and female-associated genes by red circles. Consistent with the targeted gene-expression analysis, male Off-HFD hearts showed interaction clusters associated with reactivation of fetal program, Hippo signaling, myocardial stress responses, and hypertrophic remodeling. In contrast, the female Off-HFD network showed a distinct interaction profile characterized by genes linked to cardioprotective signaling, metabolic regulation, and suppression of pathological hypertrophy.

## DISCUSSION

About 40% of women of reproductive age in the United States are obese (4). Epidemiological evidence strongly links maternal obesity to adverse cardiometabolic health in offspring, including increased disease risks across the lifetime (9). More specifically, maternal obesity is associated with cardiovascular remodeling in offspring, including increased interventricular septal thickness, and a higher prevalence of hypertension (71, 72). However, in human studies, confounding variables, including shared family lifestyle and the offspring’s own diet and activity patterns, make it difficult to distinguish prenatal programming effects from postnatal environmental influences. Animal studies offer an important advantage in this regard, as they allow controlled manipulation of pregestational, gestational, and postnatal diets and hence determination of the specific contributions of each respective exposure to offspring cardiovascular health.

In the present study, we used our mouse model of maternal diet–induced obesity to determine how the maternal environment influences offspring cardiac function, with a focus on functional, metabolic, and immune alterations in offspring hearts from weaning through adulthood. Consistent with epidemiological findings, our data revealed offspring of HFD-fed mothers to develop later-life dyslipidemia, cardiac hypertrophy, hypertension, and functional abnormalities, all despite being maintained on a regular diet after weaning and raised under controlled, low-stress conditions. Importantly, our functional data demonstrate sexually dimorphic cardiac adaptation to maternal obesity: male offspring developed hypertension, increased septal and posterior wall thickness, and cardiac fibrosis, whereas female offspring exhibited a less severe phenotype.

Sexual dimorphism in the fetal cardiac response to maternal obesity has been reported previously. An analysis of more than 2 million live singleton infants without congenital malformations in Sweden showed male sex to be associated with higher rates of cardiovascular disease than female sex (13), Similarly, a literature search of 96 articles by Matuszak and colleagues (73) identified significantly greater occurrence of coronary heart disease in males born to mothers with obesity than in females. In our study, although both male and female Off-HFD exhibited cardiac hypertrophy, male offspring showed a molecular profile more indicative of pathological cardiac stress, including increased expression of *Nppa* and *Nppb,* two cardiac hormones critical for cardiac development (74). Reactivation of the fetal gene program in the adult heart is a well-established feature of adverse cardiac remodeling, and Nppa/Nppb are widely used as markers of cardiac stress and heart failure risk (74). Distinct from Off-HFD males, female Off-HFD hearts did not show activation of these pathological markers, but instead demonstrated increased expression of genes associated with cardioprotective pathways, including *Gper1* (67), *Rasd1* (68), and *Klf15* (69). This pattern is consistent with extant evidence that biological sex specifically modifies cardiac remodeling responses, with female hearts often showing greater resistance to pathological hypertrophy and adverse remodeling (75). Activation of YAP/TAZ signaling that we observed in male, but not female, Off-HFD hearts provides a potential mechanistic link between maternal obesity and the male-specific hypertrophic transcriptional program. YAP and TAZ are downstream effectors of the Hippo pathway that regulate cardiomyocyte growth, stress adaptation, fibrosis, and cardiac remodeling. Transient YAP/TAZ activation supports compensatory hypertrophic growth, but its sustained or dysregulated activation has been implicated in pathological remodeling and progression toward heart failure (76). Together, these findings suggest that maternal obesity programs distinct cardiac molecular trajectories in male and female offspring: males develop a more pathological hypertrophy-related transcriptional profile, whereas females activate a more adaptive or cardioprotective response.

A notable finding from this study is that Off-HFD mice appear to rely heavily on lipids as an energy source. The combination of low RER, hyperlipidemia, and altered cardiac lipid and mitochondrial metabolism suggests both systemic and heart-specific metabolic inflexibility, characterized by increased fatty acid utilization, excess lipid availability, and impaired myocardial lipid handling. Importantly, human studies support the clinical relevance of this finding. Pericardial fat accumulation, similar to that observed in Off-HFD mice, has been independently associated with subclinical atrial and right ventricular dysfunction(77) as well as with increased incidence of coronary heart disease in the Multi-Ethnic Study of Atherosclerosis (78). In addition, in male participants of the Framingham Heart Study, pericardial fat was directly correlated with left ventricular mass (79). This shift toward greater lipid dependence is often accompanied by broader disturbances in whole-body lipid trafficking, including enhanced fatty acid mobilization from storage depots, increased hepatic lipid synthesis, and reduced clearance of circulating lipids (80). Hence, changes in blood lipids in the Off-HFD may reflect a compensatory increase in lipid supply to meet tissue energy demands, as fatty acids are mobilized from adipose triglyceride stores and transported to peripheral tissues for oxidation (81). In parallel, these changes may also indicate that lipid delivery exceeds tissue uptake and oxidative capacity, resulting in accumulation of circulating lipid species despite preferential lipid use at the tissue level (80); this is in keeping with the elevated blood lipid profile we observed in Off-HFD mice. Thus, increased circulating cholesterol, triglycerides, and phospholipids may not contradict preferential lipid utilization, but rather represent evidence of dysregulated lipid trafficking in which lipid supply, transport, and use are all increased but remain poorly coordinated.

Such dysregulated lipid metabolism likely led to the cardiac ectopic lipid deposition that we observed by PAI, which may have adverse functional consequences; that is, excess lipid accumulation can disrupt mitochondrial structure and function, activate oxidative stress, and impair cardiac bioenergetics. Congruent with these effects, our ultrastructural and molecular analyses revealed evidence of mitochondrial abnormalities in Off-HFD hearts. As the heart relies heavily on mitochondrial oxidative metabolism to sustain contractile function, these defects would be expected to increase cellular stress and reduce energetic efficiency, thereby contributing to maladaptive cardiac remodeling. In addition, the reduced oxygenated hemoglobin signal detected by PAI is consistent with diminished myocardial oxygen availability and/or utilization, further supporting the presence of metabolic dysfunction in Off-HFD hearts.

Our epigenetic and transcriptomic analyses provide additional support for this interpretation. Namely, we identified several genes involved in lipid metabolism that were differentially methylated in newly weaned offspring and also differentially expressed in adult offspring, including *Stard3* (82), *Sparcl1* (83), and *Copg1* (84). These findings suggest that epigenetic alterations in lipid regulatory pathways emerge early in life, before the progression of overt metabolic disease, and may persist into adulthood. Importantly, the heart does not appear to be the only organ affected by lipid metabolic dysregulation following exposure to maternal obesity; in previous studies, we showed maternal obesity to also alter lipid metabolism in the bone marrow of newly weaned offspring (19), further supporting the idea that disrupted lipid handling begins early and extends beyond classical metabolic tissues. Consistent with this concept, we have previously identified lipid metabolic abnormalities in the placenta and cord blood of infants born to obese mothers (85), suggesting that altered lipid trafficking and utilization are already present at the maternal–fetal interface and in the fetal circulation. Collectively, these observations indicate that lipid dysregulation may be a common and persistent feature of developmental programming by maternal obesity. Thus, rather than representing an isolated cardiac phenotype, altered myocardial lipid metabolism may reflect a broader systemic metabolic program established during fetal development and maintained across multiple offspring tissue types.

Beyond lipid metabolism, we also identified nineteen genes that exhibited consistent differential methylation in newly weaned (3-week) offspring and adult (6-month) offspring, eleven of which have previously been linked to cardiac development, extracellular matrix organization, cardiac hypertrophy, and hypertension. The persistence of these methylation changes from weaning into adulthood is in keeping with the developmental origins of health and disease paradigm, which proposes that exposure to an adverse intrauterine environment can leave long-lasting epigenetic marks that shape tissue function and influence later disease susceptibility (86).

Together, our findings indicate that offspring cardiac adaptation to maternal obesity is underscored by metabolic, immune, and functional changes. This in turn prompts questions regarding the timing and underlying mechanism in which the maternal environment affects fetal cardiac development. Alterations in epigenetic modifications appear to address this question at least partially, as such modifications are already known to regulate numerous critical processes in the heart (87). DNA methylation in particular is increasingly recognized as an important regulator of cardiomyocyte development, metabolic maturation, stress-response signaling, and pathological remodeling (88). Thus, the presence of stable differentially methylated genes in Off-HFD mice suggests that maternal obesity may establish a durable cardiac epigenetic memory, potentially priming the offspring heart for altered transcriptional responses, impaired metabolic adaptation, and increased vulnerability to hypertrophic remodeling later in life (89). Epigenetic modifications have already been described in the setting of maternal obesity (90); for example, a clinical study of 40 mother-infant dyads revealed strong association of maternal blood metabolites with DNA methylation patterns in fetal umbilical cord blood (91), and exposure to maternal obesity during development is associated with perturbation of global DNA methylation at CpG sites and islands in the offspring’s white adipose tissue (92).

Although the metabolic, functional, and immune perturbations described here are not immediately catastrophic in and of themselves, they may increase the vulnerability of the offspring heart to secondary nutritional, psychological, or environmental insults, thereby elevating the risk of future cardiovascular disease. Heart disease remains the leading cause of death in the United States, and many of its major risk factors, including hypertension and obesity, continue to rise at alarming rates (2). While unhealthy dietary and lifestyle choices undoubtedly contribute to cardiovascular disease risk, the present findings suggest that exposure to maternal obesity per se may predispose offspring to cardiac dysfunction independent of their own lifestyle. Defining the mechanisms that underlie this programmed susceptibility could open new opportunities to prevent cardiac dysfunction in future offspring of mothers with obesity. In particular, targeted modulation of lipid metabolism or immune-cell responses or mitochondrial function may represent promising strategies to reduce fibrosis, improve myocardial energy metabolism, and limit progression toward pathological cardiac remodeling.

The study had several limitations, DNA methylation and RNA-sequencing analyses were performed on whole-heart tissue rather than at single-cell resolution and therefore we cannot determine whether the observed epigenetic and transcriptional changes reflect cell-intrinsic alterations within specific cardiac populations. In addition, the RRBS analyses at different mouse ages were performed independently with slightly different methods, although we expect results to be reasonably comparable despite this variation in methods. Population specificity is important to consider given the immune, fibrotic, and metabolic perturbations observed in Off-HFD hearts, which likely involve coordinated responses across multiple cell types. Single-cell or single-nucleus multi-omic approaches would allow future studies to define the particular cell populations that carry persistent DNA methylation changes, determine how these changes relate to gene expression within the same cell types, and identify whether maternal obesity preferentially programs cardiomyocyte metabolic pathways, fibroblast activation, vascular dysfunction, or immune-cell remodeling. Such analyses will be important for distinguishing primary programmed effects from secondary tissue remodeling and for identifying the cellular mechanisms that drive sex-specific cardiac vulnerability in offspring exposed to maternal obesity.

In the public health context, pharmacological interventions for obesity continue to advance, but the difficulty of achieving and maintaining sustained weight loss due to frequent relapse makes prevention imperative. While lifestyle choices are well-recognized determinants of obesity and its associated comorbidities, including cardiovascular disease, findings from our study and others (93) highlight that *in utero* exposure to a maternal obesogenic diet can directly contribute to increased risk in offspring. As such, effective prevention strategies should begin prior to conception and continue across the lifespan.

## Declaration of Interests

none

## Grant Support and Acknowledgements

We thank Kim Montaniel for his assistance with mean arterial pressure measurements, Dr. Anna St. Lorenz from FUJIFILM VisualSonics for her assistance with photoacoustic imaging, and Dr. Beth Habecker for valuable discussions of our data. Financial support for this work was provided by the NIH HL170097, HL164474, HD099367, HD118370, OHSU Exploratory Research Seed Grants, and OHSU Medical Research Foundation (to A.M.), NIH/NIAMS R01AR080150 (to KN and JV), and 1S10OD038322 (to SR). The University of Cincinnati Mouse Metabolic Phenotyping Center was funded by the NIDDK (National MMPC, RRID:SCR_008997, www.mmpc.org) under the MICROMouse Program, Grants DK076169. We acknowledge the KCVI Epigenetics Consortium, OHSU Histopathology Core, the Massive Parallel Sequencing Shared Resource, and the Oregon National Primate Research Center Bioinformatics & Biostatistics Core (NIH P51 OD011092), and the OHSU Advanced Computing Center (NIH S10 OD034224).

## Author Contributions

SK, SR and AM conceived and designed the study; TDW, EAP, YJA, CB, JV and ZZ conducted the experiments and analyzed the data; SBG provided guidance and access to the tail cuff blood pressure measuring system; ShK and JR analyzed echocardiography data; SK planned flow cytometry experiments and assisted with data analysis; BD, LC, ShR, SSF, LS performed bioinformatics and biostatistics analyses of the DNA methylation and RNA-seq data; YC, SR, JL, and KN provided an access to instrumentation and analytic tools; SR and AM acquired funding; AM wrote the manuscript. All authors provided feedback and assisted with preparing the final manuscript.

## Data Availability

DNA methylation and RNA-seq data were submitted to Gene Expression Omnibus (GEO) repository, the accession number is GSE294642.

## List of Abbreviations

ATG7: Autophagy Related 7
BMI: Body Mass Index
DEG: Differentially Expressed Genes
DMR: Differentially Methylated Regions
HFD: High Fat Diet
IPA: Ingenuity Pathway Analysis
LC3: Microtubule-associated protein 1A/1B-light chain 3
Micro-CT: Micro-Computed Tomography
Off-HFD: Offspring of HFD-fed mothers
Off-RD: Offspring of RD-fed mothers
PAI: Photoacoustic Imaging
RD: Regular Diet
RER: Respiratory Exchange Ratio
RRBS: Reduced Representation Bisulfite Sequencing
STRING: Search Tool for the Retrieval of Interacting Genes/Proteins
TAZ: Transcriptional co-activator with PDZ-binding motif.
TFEB: Transcription factor EB
5’-UTR: 5’-Untranslated Regions
YAP: Yes-Associated Protein

## Notes

### Competing Interest Statement

The authors have declared no competing interest.

### Summary of Updates

New data include: Micro-CT imaging Photoacoustic imaging DNA methylation RNA-sequencing

