## Supplemental Tables for "Sex-Dependent Epigenomic and Transcriptomic Reprogramming Links Maternal Obesity to Cardiac Remodeling in Adult Offspring"

| Gene | chr | Length | % methyl  change | Sidak_Pval | Avg methyl  Off-HFD | Avg methyl  Off-RD | Gene description |
| --- | --- | --- | --- | --- | --- | --- | --- |
| H13 | 2 | 180 | -43.65 | 7.36E-10 | 41.71 | 85.37 | Minor histocompatibility antigen H13 |
| Impact | 18 | 288 | -42.68 | 6.33E-17 | 40.12 | 82.81 | Impact, RWD domain protein (Impact) |
| Snx20 | 8 | 13 | -42.13 | 2.4E-02 | 9.98 | 52.12 | Sorting nexin 20 |
| Lamtor3 | 3 | 75 | -28.36 | 4.99E-03 | 29.12 | 57.48 | Late endosomal/lysosomal adaptor, MAPK and MTOR activator 3 |
| Mindy4 | 6 | 71 | -27.28 | 9.49E-07 | 26.45 | 53.74 | Probable ubiquitin carboxyl-terminal hydrolase MINDY-4 |
| Pecam1 | 11 | 74 | -26.98 | 4.26E-04 | 19.35 | 46.34 | Platelet/endothelial cell adhesion molecule 1 |
| Kcnb1 | 2 | 18 | -26.81 | 2.22E-02 | 20.90 | 47.72 | Potassium voltage-gated channel subfamily B member 1 |
| Dock9 | 14 | 74 | -24.12 | 1.94E-04 | 31.11 | 55.24 | Dedicator of cytokinesis protein 9 isoform 5 |
| Elk3 | 10 | 21 | -22.95 | 7.83E-04 | 36.53 | 59.49 | ETS domain-containing protein Elk-3 |
| Eng | 2 | 170 | -22.49 | 9.75E-03 | 24.03 | 46.52 | Endoglin |

**Table S1**. Hypomethylated DMRs with Sidak p-value<0.05 and percentage of methylation change between Off-RD and Off-HFD >10%.

| Gene | Chr | Length | % methyl change | Sidak_Pval | Avg methyl Off-HFD | Avg methyl Off-RD | Gene description |
| --- | --- | --- | --- | --- | --- | --- | --- |
| Twf1 | 15 | 75 | 25.10 | 2.11E-03 | 42.73 | 17.62 | Twinfilin actin binding protein 1 |
| Golim4 | 3 | 74 | 24.46 | 3.08E-04 | 26.26 | 1.80 | Golgi integral membrane protein 4 |
| Atp6v1g1 | 4 | 50 | 23.18 | 3.63E-06 | 25.59 | 2.41 | ATPase, H+ transporting, lysosomal V1 subunit G1 |
| Notch1 | 2 | 74 | 17.58 | 8.98E-03 | 18.34 | 0.76 | Neurogenic locus notch homolog protein 1 |
| Adamts1 | 16 | 74 | 12.23 | 3.94E-02 | 13.40 | 1.16 | A disintegrin and metalloproteinase with thrombospondin motifs 1 |
| Gle1 | 2 | 68 | 8.89 | 1.53E-02 | 9.94 | 1.04 | Nucleoporin GLE1 |
| Cebpa | 7 | 116 | 7.82 | 8.72E-06 | 9.65 | 1.82 | CCAAT/enhancer-binding protein alpha |
| Mzf1 | 7 | 127 | 7.45 | 5.35E-03 | 9.46 | 2.01 | Myeloid zinc finger 1 |
| C2cd4b | 9 | 54 | 6.98 | 1.00E-02 | 9.67 | 2.69 | C2 calcium-dependent domain containing 4B |
| Cadm1 | 9 | 203 | 5.21 | 1.79E-03 | 5.46 | 0.24 | Cell adhesion molecule 1 |

**Table S2.** Hypermethylated DMRs with Sidak p-value<0.05 and percentage of methylation change between Off-RD and Off-HFD >10%.

| Symbol | Description | Relation to CVD |
| --- | --- | --- |
| ARHGAP22 | Rho GTPase Activating Protein 22 | Variations in blood DNA methylation near the gene that are significantly associated with the incidence of **coronary heart disease**(1) |
| Asap2 | ArfGAP with SH3 domain, ankyrin repeat and PH domain 2 | Differentially methylated in hypertension (2) |
| CMIP | C-Maf Inducing Protein | Genetic variations in CMIP are linked to **dyslipidemia and fatty liver** **(3, 4)** |
| ****CTBP1**** | C-terminal binding protein 1 | Cardiomyocyte hypertrophy(5) |
| ****Elk3**** | ETS transcription factor | Highly expressed in heart tissue during conditions like dilated cardiomyopathy or hypertrophic cardiomyopathy(6). |
| Eng | Endoglin | Hypertension, hypertrophic cardiomyopathy(7). |
| ****Esam**** | Endothelial cell‐selective adhesion molecule | Deficiency results in reduced coronary microvascular density (8). |
| ****Kcnb1**** | KV2.1 is a voltage-gated delayed rectifier potassium channel | Mutations associated with Arrythmia and Long QT syndrome (9). |
| Mindy4 | MINDY lysine 48 deubiquitinase 4 | Hypertension, high triglycerides (10). |
| Npc1 | Niemann-Pick C1 | Regulates intracellular/lysosomal cholesterol trafficking (11) |
| Ppcdc | Phosphopantothenoylcysteine decarboxylase | Regulates CoA biosynthesis essential for fatty acid synthesis and β-oxidation (12). |
| PCSK6 | Proprotein convertase subtilisin/kexin 6 | Dysregulation was linked to atherosclerosis, cardiomyocyte senescence (13) hypertension, atrial fibrillation, and myocardial infarction (14). |
| SH3PXD2A | SH3 and PX domains 2A. | Extracellular matrix organization(15), coronary artery disease(16), cardiac conduction block(17) |
| ****SVIL**** | Actin-binding protein **supervillin** | Recognized as a novel disease gene for **hypertrophic cardiomyopathy (HCM).** |
| TRAPPC9 | Trafficking Protein Particle Complex Subunit 9 | Regulates lipid droplet formation (18) |

**Table S3.** Comparison of two RRBS datasets from 3-week-old and 6-month-old Off-HFD mice revealed DMRs of genes associated with cardiometabolic diseases.

| **Symbol** | **Description** | **DMR**  **6 mo** | **DEG**  **6mo** |
| --- | --- | --- | --- |
| Abracl | ABRA C-terminal like | ↓ | ↓ |
| Agtpbp1 | ATP/GTP binding carboxypeptidase 1 | ↓ | ↓ |
| Akap7 | A-Kinase Anchoring Protein 7 | ↑ | ↓ |
| Amotl2 | Angiomotin-like protein 2 | ↑ | ↑ |
| Aopep | Aminopeptidase O | ↑ | ↓ |
| Arhgap6 | Rho GTPase activating protein 6 | ↑ | ↓ |
| Arl6ip1 | ADP-Ribosylation Factor-Like GTPase 6 Interacting Protein 1 | ↓ | ↑ |
| Armc5 | armadillo repeat containing 5 | ↓ | ↑ |
| Bad | BCL2-associated agonist of cell death | ↑ | ↓ |
| Bdkrb2 | Bradykinin receptor, beta 2 | ↓ | ↑ |
| Brf1 | BRF1, RNA polymerase III transcription initiation factor 90 kDa subunit | ↑ | ↓ |
| Btg2 | BTG anti-proliferation factor 2 | ↑ | ↑ |
| Cacnb4 | Calcium channel, voltage-dependent, beta 4 subunit | ↑ | ↓ |
| Cbx4 | Chromobox 4 | ↑ | ↑ |
| Cbx8 | Chromobox 8 | ↓ | ↑ |
| Ccne2 | Cyclin E2 | ↑ | ↑ |
| Cdc42bpa | CDC42 binding protein kinase alpha | ↓ | ↑ |
| Cdca4 | Cell division cycle associated 4 | ↑ | ↑ |
| Cdca7l | Cell division cycle associated 7 like | ↓ | ↑ |
| Cemip2 | Cell migration inducing hyaluronidase 2 | ↓ | ↑ |
| Cenpm | Centromere protein M | ↓ | ↑ |
| Cpe | Carboxypeptidase E | ↑ | ↓ |
| Cspp1 | Centrosome and spindle pole associated protein 1 | ↓ | ↓ |
| Cxcl12 | C-X-C motif chemokine ligand 12 | ↑ | ↑ |
| Dcaf13 | DDB1 and CUL4 associated factor 13 | ↑ | ↓ |
| Dgkb | Diacylglycerol kinase, beta | ↑ | ↓ |
| Dnajc10 | DnaJ heat shock protein family (Hsp40) member C10 | ↓ | ↑ |
| Dnm1 | Dynamin 1 | ↓ | ↓ |
| E230016M11Rik | RIKEN cDNA E230016M11 gene | ↓ | ↓ |
| E530011L22Rik | RIKEN cDNA E530011L22 gene | ↓ | ↓ |
| Ets2 | E26 avian leukemia oncogene 2, 3' domain | ↑ | ↑ |
| Fbxo34 | F-box protein 34 | ↓ | ↑ |
| Gcsh | Glycine cleavage system protein H (aminomethyl carrier) | ↓ | ↓ |
| Ggta1 | Glycoprotein galactosyltransferase alpha 1, 3 | ↓ | ↑ |
| Gpatch2l | G patch domain containing 2 like | ↑ | ↓ |
| Gprc5b | G protein-coupled receptor, family C, group 5, member B | ↑ | ↑ |
| H4c9 | H4 clustered histone 9 | ↓ | ↓ |
| Hacd1 | 3-hydroxyacyl-CoA dehydratase 1 | ↑ | ↓ |
| Hdac9 | Histone deacetylase 9 | ↑ | ↑ |
| Inpp5b | inositol polyphosphate-5-phosphatase B | ↓ | ↑ |
| Iqsec1 | IQ motif and Sec7 domain 1 | ↑ | ↓ |
| Katnal1 | Katanin p60 subunit A-like 1 | ↑ | ↑ |
| Kbtbd7 | Kelch repeat and BTB (POZ) domain containing 7 | ↑ | ↑ |
| Kcnd3 | Potassium voltage-gated channel, Shal-related family, member 3 | ↑ | ↓ |
| Kcnh2 | Potassium voltage-gated channel, subfamily H (eag-related), member 2 | ↑ | ↓ |
| Kctd2 | Potassium channel tetramerization domain containing 2 | ↑ | ↑ |
| Kif1a | Kinesin family member 1A | ↑ | ↓ |
| Kifc5b | Kinesin family member C5B | ↑ | ↑ |
| Lamb1 | Laminin B1 | ↑ | ↑ |
| Lhx6 | LIM homeobox protein 6 | ↑ | ↑ |
| Lmnb1 | Lamin B1 | ↓ | ↑ |
| Lonrf1 | LON peptidase N-terminal domain and ring finger 1 | ↑ | ↑ |
| Lrrc28 | Leucine rich repeat containing 28 | ↑ | ↑ |
| Mex3c | Mex3 RNA binding family member C | ↓ | ↑ |
| Mfn2 | Mitofusin 2 | ↑ | ↓ |
| Mgat4a | mannoside acetylglucosaminyltransferase 4, isoenzyme A | ↑ | ↑ |
| Mphosph10 | M-phase phosphoprotein 10 (U3 small nucleolar ribonucleoprotein) | ↓ | ↑ |
| Mrpl37 | Mitochondrial ribosomal protein L37 | ↑ | ↓ |
| Mrps35 | Mitochondrial ribosomal protein S35 | ↓ | ↓ |
| Notch2 | Notch 2 | ↓ | ↓ |
| Pard6g | Par-6 family cell polarity regulator gamma | ↓ | ↓ |
| Pde8a | Phosphodiesterase 8A | ↓ | ↑ |
| Pdgfd | platelet-derived growth factor, D polypeptide | ↑ | ↑ |
| Plekha7 | Pleckstrin homology domain containing, family A member 7 | ↑ | ↑ |
| Plgrkt | plasminogen receptor, C-terminal lysine transmembrane protein | ↓ | ↓ |
| Polr2m | Polymerase (RNA) II (DNA directed) polypeptide M | ↑ | ↓ |
| Psmd1 | proteasome (prosome, macropain) 26S subunit, non-ATPase, 1 | ↓ | ↑ |
| Ptpra | Protein tyrosine phosphatase receptor type A | ↑ | ↑ |
| Rapgef5 | Rap guanine nucleotide exchange factor (GEF) 5 | ↑ | ↑ |
| Rasgrp2 | RAS, guanyl releasing protein 2 | ↑ | ↓ |
| Rcsd1 | RCSD domain containing 1 | ↓ | ↑ |
| Rreb1 | Ras responsive element binding protein 1 | ↓ | ↑ |
| Rtel1 | Regulator of telomere elongation helicase 1 | ↑ | ↓ |
| Samd12 | Sterile alpha motif domain containing 12 | ↓ | ↑ |
| Scn4b | Sodium channel, type IV, beta | ↓ | ↑ |
| Sgms1 | Sphingomyelin synthase 1 | ↓ | ↑ |
| Siva1 | SIVA1, apoptosis-inducing factor | ↑ | ↓ |
| Slc46a3 | Solute carrier family 46, member 3 | ↑ | ↑ |
| Sstr3 | Somatostatin receptor 3 | ↓ | ↓ |
| Sstr4 | Somatostatin receptor 4 | ↑ | ↑ |
| Stard3 | StAR related lipid transfer domain containing 3 | ↑ | ↓ |
| Tbx15 | T-box 15 | ↓ | ↑ |
| Thap4 | THAP domain containing 4 | ↓ | ↓ |
| Tm7sf3 | Transmembrane 7 superfamily member 3 | ↑ | ↑ |
| Tm9sf1 | Transmembrane 9 superfamily member 1 | ↓ | ↑ |
| Tmcc1 | Transmembrane and coiled coil domains 1 | ↓ | ↑ |
| Tpm2 | Tropomyosin 2, beta | ↓ | ↓ |
| Ttll1 | Tubulin tyrosine ligase-like 1 | ↑ | ↓ |
| Uck2 | Uridine-cytidine kinase 2 | ↑ | ↑ |
| Uhmk1 | U2AF homology motif (UHM) kinase 1 | ↓ | ↑ |
| Usp36 | Ubiquitin specific peptidase 36 | ↑ | ↓ |
| Wdr6 | WD repeat domain 6 | ↑ | ↑ |
| Whrn | Whirlin | ↑ | ↓ |
| Ybx2 | Y box protein 2 | ↓ | ↓ |
| Zbtb34 | Zinc finger and BTB domain containing 34 | ↑ | ↑ |
| Zfp84 | zinc finger protein 84 | ↓ | ↑ |

**Table S3.** Genes showing concurrent changes in DNA methylation and gene expression in 6-month-old male and female Off-HFD hearts compared with Off-RD controls.

| Males | | | | Females | | | |
| --- | --- | --- | --- | --- | --- | --- | --- |
| Symbol | **Experim. FC** | **Z-score** | **P-value** | **Symbol** | **Experim. FC** | **Z-score** | **P-value** |
| Activated |  |  |  | **Activated** | | | |
| TCF7L2 |  | 2.06 | 5.9E-08 | **NFAT5** |  | 2.51 | 1.65E-05 |
| TCF7L1 |  | 2.16 | 1.02E-05 | **FOXO3** |  | 2.08 | 2.98E-03 |
| ATF4 |  | 2.43 | 5.98E-04 | **STAT4** |  | 2.30 | 7.08E-03 |
| PPARGC1B |  | 2.45 | 1.91E-03 | **PBX3** |  | 2.65 | 2.07E-03 |
| KLF1 |  | 2.62 | 3.24E-03 | **SPIB** |  | 2.84 | 6.67E-03 |
| SMAD3 | 5.45 | 2.26 | 1.41E-03 | **ARNT** |  | 2.39 | 3.22E-02 |
| GLIS3 | 5.31 | 2.18 | 4.04E-03 | **miR-338-3p** |  | 2.02 | 1.05E-02 |
| IRF8 |  | 2.01 | 3.15E-03 | **MEIS1** | 1.53 | 2.81 | 2.42E-02 |
| SREBF1 |  | 2.42 | 4.29E-02 | **TRIM28** |  | 2.32 | 2.46E-02 |
| STAT4 |  | 3.23 | 1.44E-02 |  |  |  |  |
| Inhibited | | | | **Inhibited** | | | |
| mir-8 |  | -2.55 | 2.92E-05 | **PAX6** | 1.83 | -2.39 | 1.25E-07 |
| FOXA1 |  | -2.07 | 1.64E-05 | **LMO2** | 3.08 | -2.14 | 1.18E-06 |
| miR-124-3p |  | -2.02 | 1.54E-04 | **mir-210** |  | -2.25 | 1.47E-05 |
| miR-29b-3p |  | -2.44 | 2.36E-04 | **EOMES** |  | -2.04 | 9.71E-04 |
| miR-1-3p |  | -3.14 | 1.45E-04 | **MYCN** |  | -3.00 | 9.71E-04 |
| miR-27a-3p |  | -2.90 | 1.06E-03 | **Rhox5** |  | -2.14 | 4.36E-03 |
| miR-30c-5p |  | -2.35 | 3.06E-03 | **mir-30** | -5.49 | -2.10 | 1.06E-03 |
| let-7a-5p |  | -2.16 | 6.92E-03 | **mir-135** |  | -2.37 | 1.57E-03 |
| mir-140 |  | -2.12 | 2.28E-03 | **ETV4** | -6.31 | -2.12 | 1.72E-03 |
| mir-9 |  | -2.01 | 4.69E-03 | **HMGA1** |  | -3.12 | 9.86E-03 |
| miR-34a-5p |  | -2.01 | 4.39E-03 | **BMAL1** |  | -2.49 | 1.26E-02 |

**Table S4.** Analysis of upstream regulators affected by exposure to maternal obesity. List of selected upstream regulators derived from Ingenuity pathway analysis of RNA-Seq data. We used the IPA regulation z-score algorithm to identify upstream regulators that are expected to be either active or inhibited in male and female Off-HFD (positive or negative z-score that higher than 2.0 respectively) according to our RNA-seq data.

| Males | | |
| --- | --- | --- |
|  | **Z-scores** | |
| Upstream Regulators | **RRBS** | **RNA-seq** |
| Activated | | |
| TNF | 2.98 | 3.23 |
| TLR4 | 2.60 | 2.77 |
| MAP3K8 | 2.53 | 2.69 |
| IFNA4 | 2.45 | 2.63 |
| MYD88 | 2.53 | 2.37 |
| NFkB | 2.46 | 2.26 |
| PTPRJ | 2.11 | 2.24 |
| E2F | 2.59 | 2.20 |
| IL17A | 2.18 | 2.14 |
| Inhibited | | |
| NOTCH3 | -2.94 | -2.56 |
| miR-30c-5p | -2.35 | -1.61 |
| ADAM12 | -2.10 | -2.24 |
| spermine | -2.00 | -2.59 |
| PNPT1 | -2.63 | -2.61 |
| Females | | |
| Activated | | |
| NFAT5 | 2.51 | 3.15 |
| PRKAA1 | 2.75 | 2.63 |
| APOE | 2.17 | 2.21 |
| PRKAA2 | 2.00 | 2.18 |
| aspirin | 2.38 | 2.92 |
| Inhibited | | |
| IFNG | -2.21 | -2.87 |
| CD38 | -2.09 | -1.94 |
| MYD88 | -2.00 | -2.00 |
| IL5 | -3.16 | -2.00 |
| F2 | -2.06 | -2.00 |
| SRGN | -2.41 | -2.00 |
| CCL20 | -2.19 | -2.00 |
| CTNNB1 | -2.05 | -2.05 |
| ERK | -2.11 | -2.19 |
| CSF2 | -3.45 | -2.24 |
| IL4 | -2.24 | -2.30 |
| TNF | -3.02 | -2.34 |
| PI3K | -2.73 | -2.41 |
| STING1 | 2.59 | -2.45 |
| calcitriol | 2.25 | -2.67 |

**Table S5**. Integration of RRBS and RNA-seq from 6-month-old Off-HFD in the context of regulatory molecules identified by IPA regulation z-score algorithm.

| **Symbol** | **Description** | **DMR**  **3 wks** | **DEG**  **6mo** | **Relation to CVD** |
| --- | --- | --- | --- | --- |
| **ASPH** | Aspartate beta-hydroxylase | ↑ | ↓ | Mutations are linked to CVD, including valvular damage, cardiac failure, and sudden cardiac death (19) |
| **CHML** | CHM like Rab escort protein | ↑ | ↑ | Vesicle-mediated transport, regulation of mTOR function(20) |
| **CMIP** | C-Maf inducing protein | ↓ | ↑ | Negative regulator of T-cell signaling (21) |
| **CRIM1** | Cysteine rich transmembrane BMP regulator 1 | ↓ | ↑ | Increased levels have been observed in heart failure, correlating with high levels of pro-fibrotic markers (22) |
| **COPG1** | COPI coat complex subunit gamma 1 | ↓ | ↑ | Involved in lipid homeostasis by regulating perilipin presence at the lipid droplet surface (23) |
| **GOLIM4** | Golgi integral membrane protein 4 | ↑ | ↑ | Involved in cardio-metabolic diseases (24, 25). |
| **NCOA1** | Nuclear receptor coactivator 1 | ↑ | ↓ | Associated with obesity(26), myocardial dysfunction(27) |
| **GTF3C2** | General transcription factor IIIC subunit 2 | ↑ | ↓ | Associated with metabolic syndrome (28) |
| **GPATCH2L** | G-patch domain containing 2 like | ↑ | ↓ | Involved in RNA splicing and ribosome biogenesis  (29) |
| **JMJD6** | Jumonji domain containing 6, arginine demethylase and lysine hydroxylase | ↑ | ↓ | Epigenetic regulator crucial for heart development; prevents cardiac hypertrophy and heart failure by demethylating the p65 subunit of NFkB and is essential for angiogenesis and vascular development (30) |
| **MAZ** | MYC associated zinc finger protein | ↑ | ↓ | Lipid metabolism related immune marker (31) |
| **MCM8** | Mini chromosome maintenance 8 homologous recombination repair factor | ↑ | ↑ | Facilitating mitophagy (32) |
| **MRPL51** | Mitochondrial ribosomal protein L51 | ↑ | ↓ | Vital for oxidative phosphorylation, and its dysfunction is linked to mitochondrial metabolic disorders (33) |
| **PLAGL1** | PLAG1 like zinc finger 1 | ↓ | ↑ | Regulates cell proliferation and cardiac fibroblast viability(34) |
| **PTPRA** | Protein tyrosine phosphatase (PTP)receptor type A | ↑ | ↑ | Plays a crucial role in regulating cardiac hypertrophy (35) |
| **SNRPN** | Small nuclear ribonucleoprotein polypeptide N | ↓ | ↓ | Involved in cardiac development, and pathogenesis of congenital heart diseases (36) |
| **SPARCL1** | SPARC like 1 | ↓ | ↑ | Atherosclerosis(37), cholesterol and triglyceride metabolism(38) |
| **ST3GAL1** | ST3 beta-galactoside alpha-2,3-sialyltransferase 1 | ↑ | ↑ | Involved in adult onset of dilated cardiomyopathy (39) |
| **STARD3** | StAR related lipid transfer domain containing 3 | ↓ | ↑ | **Cholesterol metabolism** **(40).** |
| **PPP1R15B** | Protein phosphatase 1 regulatory subunit 15B | ↓ | ↑ | Involved in integrated stress response and progression of heart failure (41) |
| **ZFHX3** | Zinc finger homeobox 3 | ↓ | ↓ | Atrial fibrillation (42) |

**Table S6.** List and relation to cardiovascular disease of genes with differentially methylated regions at 3 weeks of age and differential expression at 6 months of age. The directions of changes are shown.
