## Supplemental Methods and Figures for "Sex-Dependent Epigenomic and Transcriptomic Reprogramming Links Maternal Obesity to Cardiac Remodeling in Adult Offspring"

**RRBS method** **and analysis**. RRBS analysis of the two age groups was performed independently by different groups, therefore methods differ slightly between analyses.

*Newly weaned mouse hearts.* DNA methylation was evaluated using RRBS (1), a genome-wide approach that allows the capture of key regulatory regions including promoters, CpG islands, and CpG island shores. RRBS libraries were generated by the Knight Cardiovascular Institute Epigenetics core for 3-week-old mice using established methods (2, 3) and by Novogene (Sacramento, CA) for 6-month-old mice. Briefly, genomic DNA from left ventricle samples were digested overnight with *Msp*I (New England Biolabs) to produce sticky ends starting with a CpG. Libraries were then prepared using the NEXTflex Bisulfite-Seq Kit (BioScientific, Avondale, AZ) and bisulfite conversion was performed with the EZ DNA Methylation-Gold Kit (Zymo Research, Irvine, California). For DNA methylation in left ventricles from 3-week old mice, libraries were multiplexed and sequenced on a NovaSeq 6000 at the OHSU Massively Parallel Sequencing Shared Resource to obtain roughly 40 million reads/library. Sequencing data were downloaded and analyzed as described in Carbone et al. (2). Briefly, after evaluation with FastQC (Andrews, 2010; http://www.bioinformatics.babraham.ac.uk/projects/fastqc), read alignment to the mouse reference genome (mm10) and methylation calling on every covered cytosine were performed with Bismark (4). Coverage files from the Bismark methylation extractor were used to obtain the coverage and methylation rate of each covered CpG. In DNA methylation of 6-month-old mice hearts, raw FASTQ sequence files were processed with our standard in-house QC and alignment protocol. Briefly, paired readsets were preprocessed with Trimmomatic and Cutadapt and aligned to the GRCm39 reference genome using Bismark to map bisulfite converted reads and determine cytosine methylation status. A methylation rate table was constructed to include only CpGs with minimum 10X coverage and found on canonical chromosomes (1-19 and X,Y). For differential analysis, the table was filtered to include CpGs with at least 10X coverage in at least two replicates per group of maternal diet (HFD or RD) and fetal sex; this subset comprised 421,177 CpGs. Limma (5) was used for differentially methylated cytosine (DMC) analysis. To stabilize variance and improve normality, methylation percentages were transformed using the **arcsin square root transformation**. A linear model was fitted with sample groups and sex as covariates (no interaction), and empirical Bayes moderation was applied to estimate DMCs. Statistical significance was determined by **moderated *t*-test**, and multiple testing correction was performed using the **Benjamini-Hochberg false discovery rate (FDR)** method. DMCs were grouped into differentially methylated regions (DMRs) with Comb-p (6), which seeds on CpGs from the DMC analysis with a p-value < 0.05 and extends the region as long as it finds another CpG with a pvalue < 0.05 within 300bp.

*Adult mouse hearts.* Methods as above until construction of the methylation rate table. In these samples, the methylation rate table was constructed to include only CpGs with minimum 10X coverage and found on canonical chromosomes (1-19 and X,Y). For differentially methylated cytosine (DMC) analysis, the table was filtered to include CpGs with at least 10X coverage in at least 70% of samples. Methylation percentages were transformed using the base 2 logit transformation and limma (27) was used to fit linear models containing outcome methylation level (M-value) with predictor variables diet group, sex, and interaction; empirical Bayes moderation was applied to estimate DMCs, and multiple testing correction was performed using the **Benjamini-Hochberg false discovery rate (FDR)** method. Differentially methylated regions (DMRs) were identified from the limma results using DMRcate (7-9) calculate combined (Fisher’s) p-values of DMCs, using DMCs with FDR p < 0.25 as input.

**RNA-sequencing analysis**.

Differential expression analysis was performed by the ONPRC Bioinformatics & Biostatistics Core. The quality of the raw sequencing files was evaluated using FastQC (Andrews S. (2010). FastQC: a quality control tool for high throughput sequence data) ccombined with MultiQC (10) (http://multiqc.info/). Trimmomatic (11) was used to remove any remaining Illumina adapters. Reads were aligned to Ensembl’s Mus_musculus. GRCm39 genome along with its corresponding annotation, release 115. The program STAR (12) (v2.7.11b) was used to align the reads to the genome. STAR has been shown to perform well compared to other RNA-seq aligners (13). Since STAR utilizes the gene annotation file, it also calculated the number of reads aligned to each gene. RNA-SeQC (14) and another round of MultiQC were utilized to ensure alignments were of sufficient quality. Gene-level raw counts were filtered to remove genes with extremely low counts in many samples following the published guidelines (15), normalized using the trimmed mean of M-values method (TMM) (16), and transformed to log-counts per million with associated observational precision weights using the voom (17) method. Gene-wise linear models with primary variables diet group, sex, and sex:diet group interaction, along with adjustment variable mean per base coverage, were employed for differential expression analyses using limma with empirical Bayes moderation (18) and false discovery rate (FDR) adjustment (19).

**Gene Ontology and pathway analysis**.

RRBS data: *Newly weaned mouse hearts.* Enrichment analysis of Gene Ontology (release date 20210101) terms was performed for genes associated with differentially methylated CpGs (DMCs) and differentially methylated regions (DMRs) using Panther (release date 2020/07/28). Hypo- and hypermethylated regions were jointly analyzed for overrepresentation using Fischer’s exact test with FDR correction (FDR<0.05), with fold enrichment of over- and underrepresented pathways reported relative to the whole mouse genome (which was used as the reference list) (33). Pathway analysis of DMR genes was conducted using STRING (Version 12.0).

RRBS data: *Adult mouse hearts.* Enrichment analysis of Gene Ontology (release date 2025) terms and transcription factors (Transcription_Factor_PPIs) was performed for differentially methylated regions (DMRs) using enrichR (R package version 3.4, <https://CRAN.R-project.org/package=enrichR). Integrative pathway analysis of DMCs and gene expression was conducted using Ingenuity Pathway Analysis (Qiagen, Redwood City, CA).

**SUPPLEMENTAL FIGURE LEGENDS**.

**Figure S1.** Western blots for expression of subunits of the mitochondrial electron transport chain in male and female Off-RD and Off-HFD, with Vinculin as loading control. Quantification data for complex I (**A**), II (**B**), III (**C**), IV (**D**), and V (**E**). Data were analyzed by two-way ANOVA followed by *t*-test, and are represented as box and whiskers with minimum and maximum and individual values with lines at mean. N=8/group of maternal diet; *p*-values are shown.

**Figure S2.** Western blots for autophagy markers TFEB (**A**), p62 (**B**), Rubicon (**C**), ATG7 (**D**), and LC3-II (**E**) in the hearts of 4-month-old Off-RD and Off-HFD. N=4–12/sex/group of maternal diet. Data are represented as individual values with lines at mean with SEM; *, *p* < 0.05 in females vs. males within each group.

**Figure S3**. Flow cytometry gating strategy for quantifying immune cell populations in myocardium from Off-RD and Off-HFD.

**Figure S4.** Principal component analysis (PCA) of differentially methylated regions, performed on 376,326 CpGs found in at least 2 replicates per 4 groups. This plot shows the female Off-RD clustering closely with the female Off-HFD, and similar clustering is apparent with males.

**Figure S5.** Multidimensional scaling (MDS) plots showing distinct separation by sex (**A**) and less-distinct separation by a group of maternal diet (**B**) with no major outliers.

**Figure S6**. **A-B,** Manhattan plots for DNA methylation in 6-month-old male (**A**) and female (**B**) Off-HFD. Chromosomal position is depicted on the x-axis, - log(*p*-value) on the y-axis. The red line indicates nominal *p*-values<1x10^-14^. **C-D,** Stacked histogram of DNA methylation distribution in male (**C**) and female (**D**) Off-HFD showing the count of CpGs located in CpG islands (red), open sea (green), and shores (blue). **E-F**, Distribution of differentially methylated CpG sites in male (**E**) and female (**F**) Off-HFD.

**Figure S7**. **Common differentially expressed genes (DEGs) affected by maternal obesity in male and female Off-HFD versus Off-RD mice.** **A,** Venn diagram showing the total number of DEGs identified in males and females, including 19 DEGs shared between sexes. **B,** Heatmap showing log2 fold-change values for the shared DEGs. All shared DEGs exhibited opposite sex-dependent patterns of regulation in response to maternal obesity.

**Figure S8**. Significant modules from transcription factor protein–protein interaction (PPI) network enrichment analysis in male (**A**) and female (**B**) Off-HFD mice.

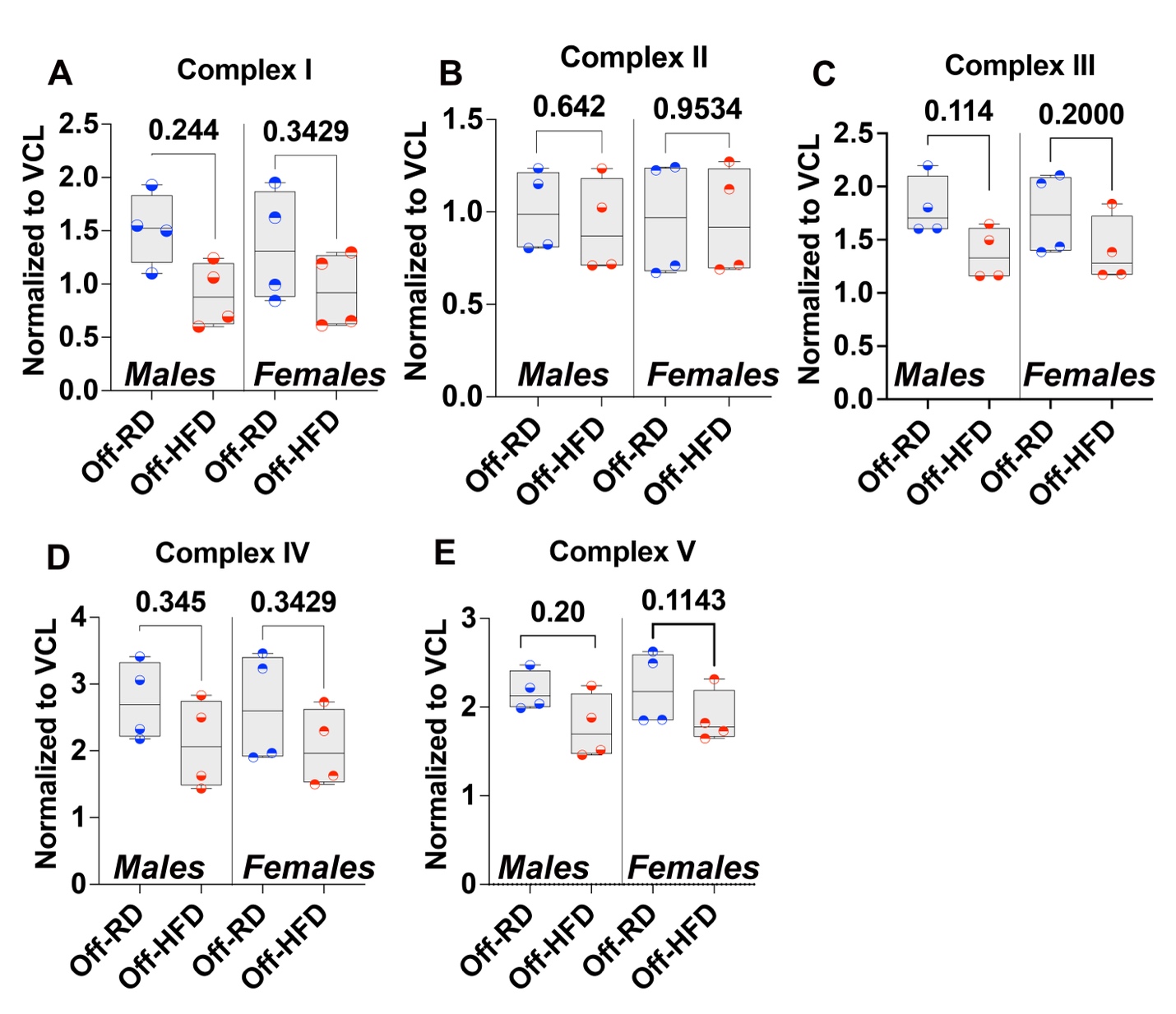

**Figure S1.** Western blots for expression of subunits of the mitochondrial electron transport chain in male and female Off-RD and Off-HFD, with Vinculin as loading control. Quantification data for complex I (**A**), II (**B**), III (**C**), IV (**D**), and V (**E**). Data were analyzed by two-way ANOVA followed by *t*-test, and are represented as box and whiskers with minimum and maximum and individual values with lines at mean. N=8/group of maternal diet; *p*-values are shown.

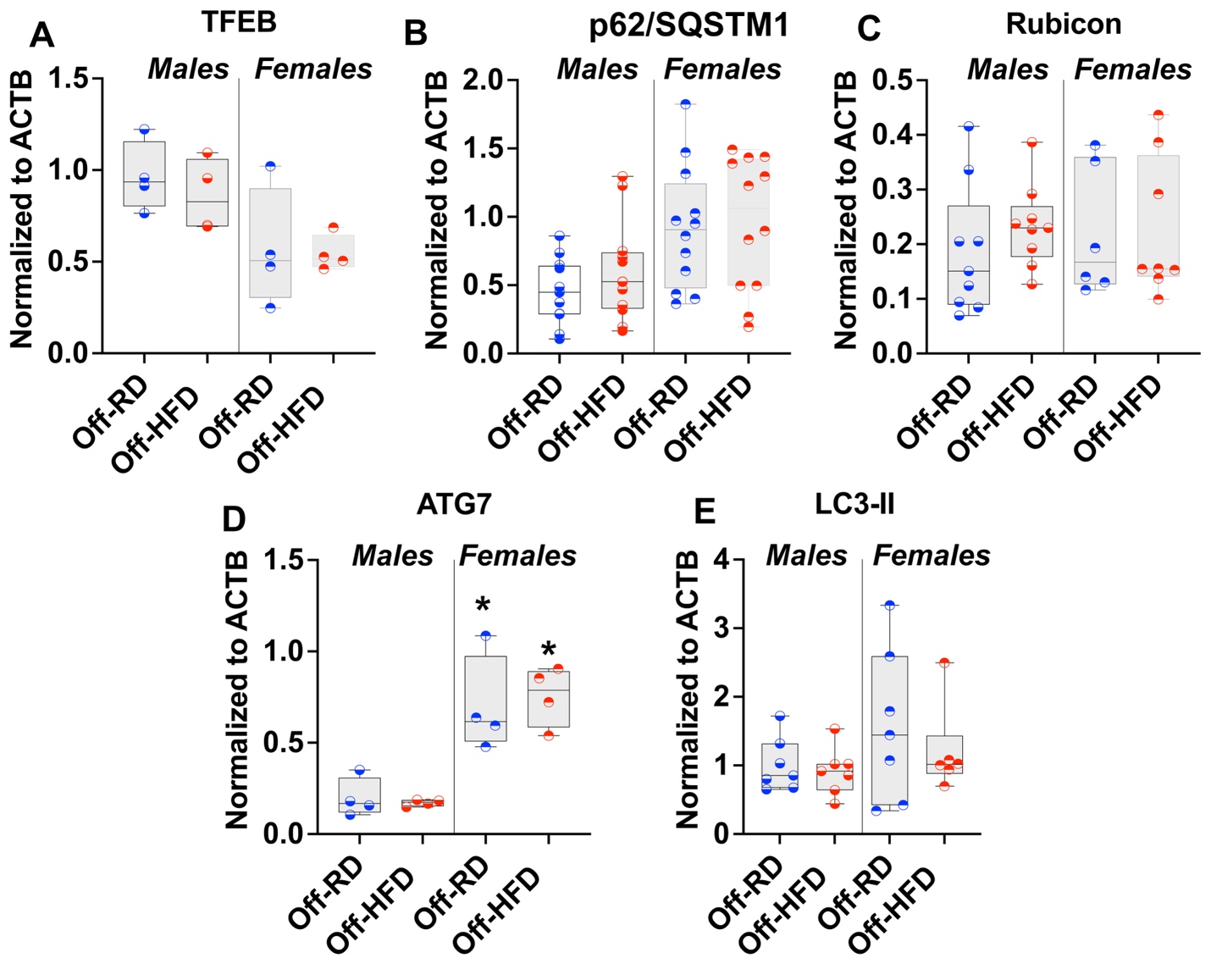

**Figure S2.** Western blots for autophagy markers TFEB (**A**), p62 (**B**), Rubicon (**C**), ATG7 (**D**), and LC3-II (**E**) in the hearts of 4-month-old Off-RD and Off-HFD. N=4–12/sex/group of maternal diet. Data are represented as individual values with lines at mean with SEM; *, *p* < 0.05 in females vs. males within each group.

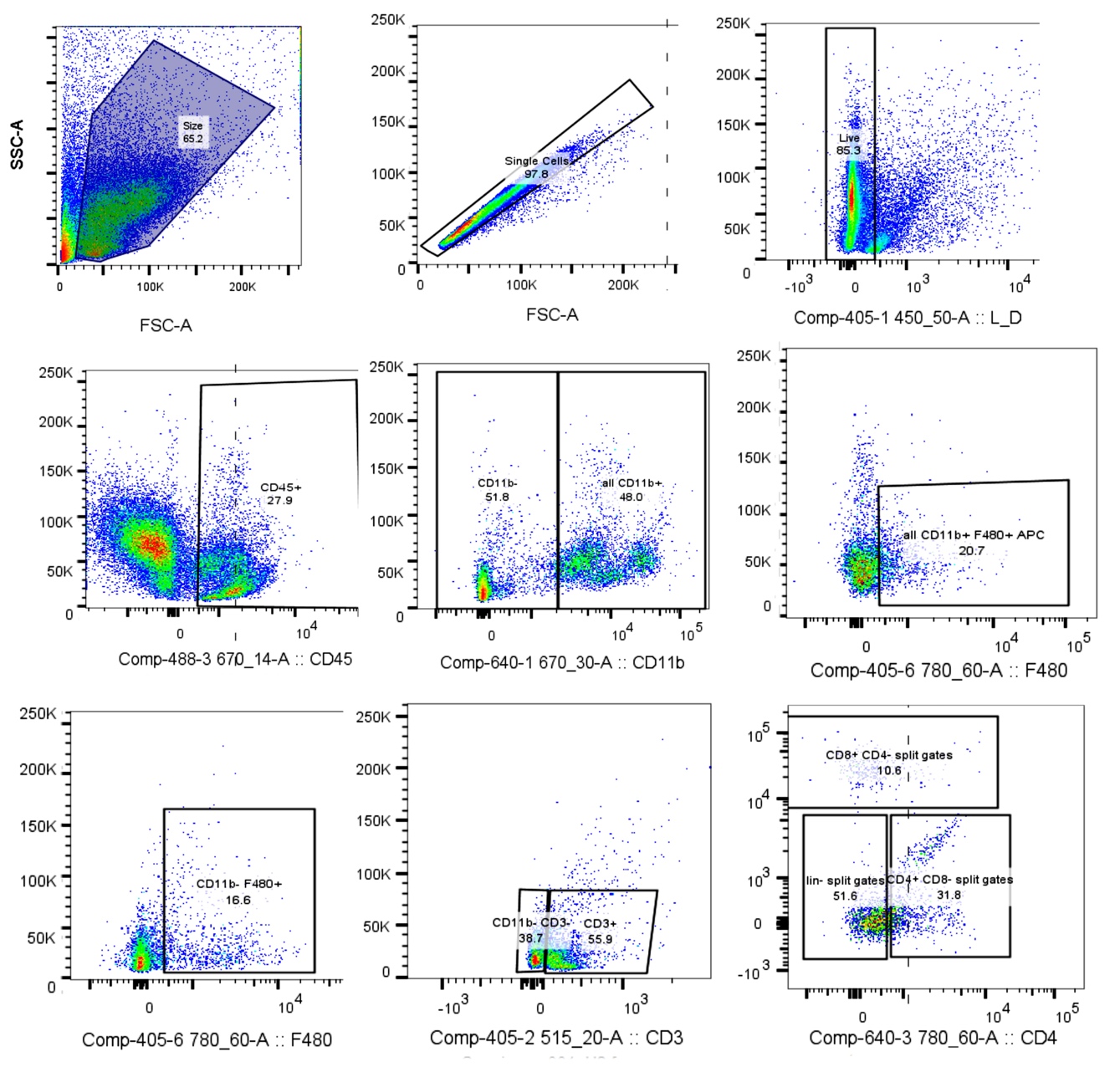

**Figure S3**. Flow cytometry gating strategy for quantifying immune cell populations in myocardium from Off-RD and Off-HFD.

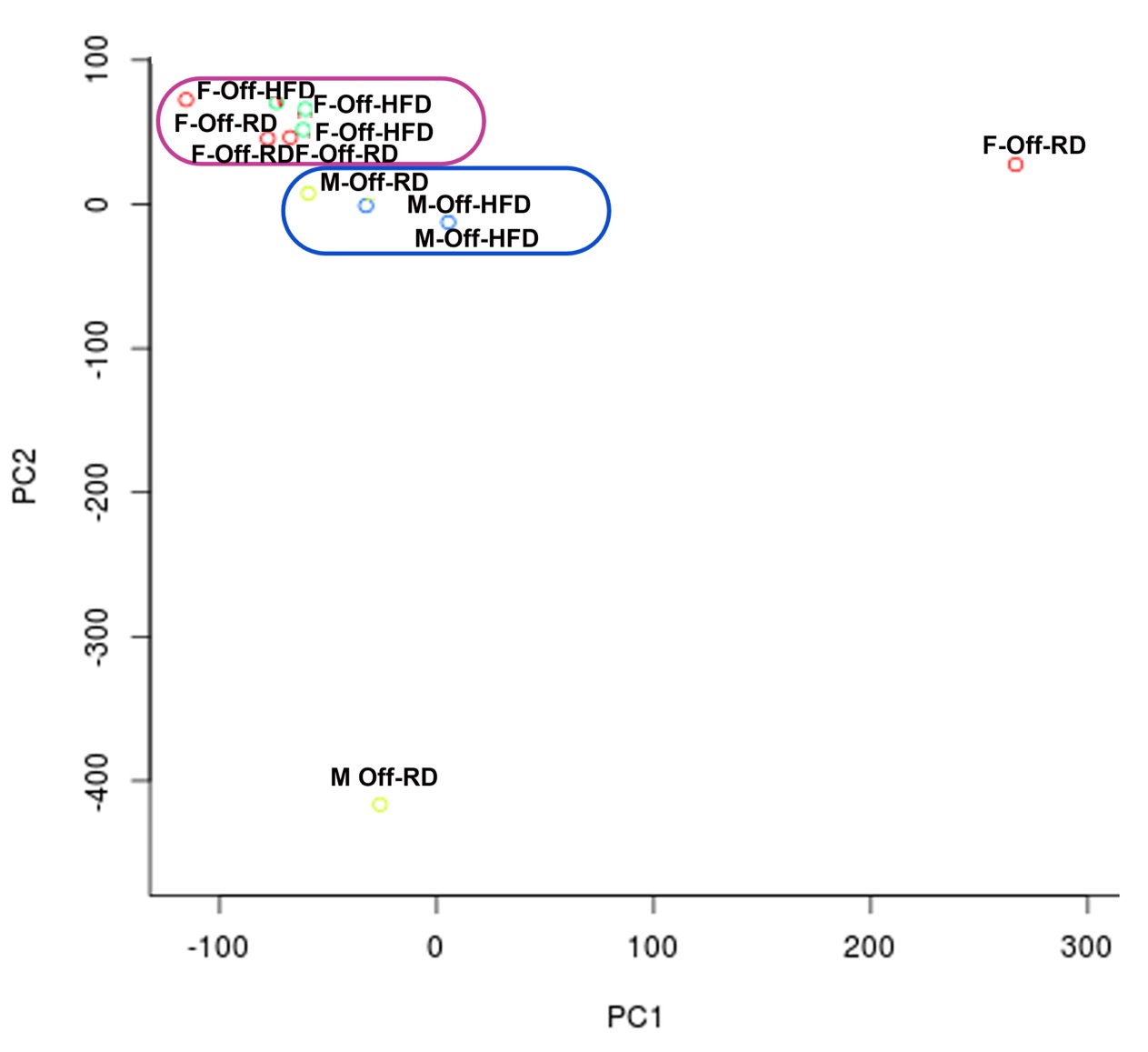

**Figure S4.** Principal component analysis (PCA) of differentially methylated regions, performed on 376,326 CpGs found in at least 2 replicates per 4 groups. This plot shows the female Off-RD clustering closely with the female Off-HFD, and similar clustering is apparent with males.

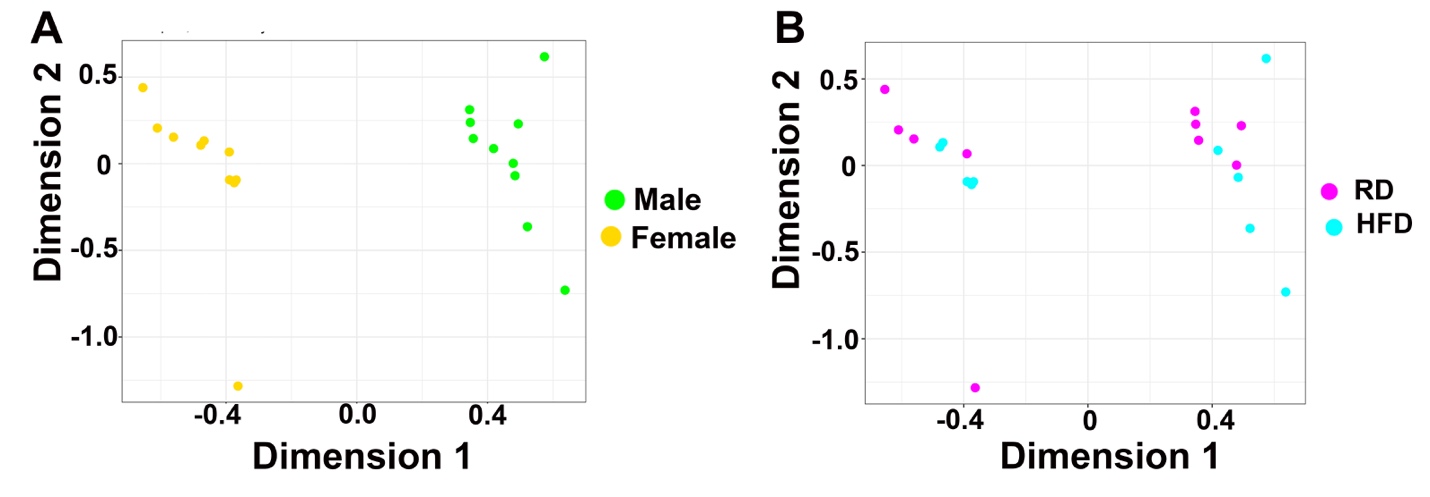

**Figure S5.** Multidimensional scaling (MDS) plots showing distinct separation by sex (**A**) and less-distinct separation by a group of maternal diet (**B**) with no major outliers.

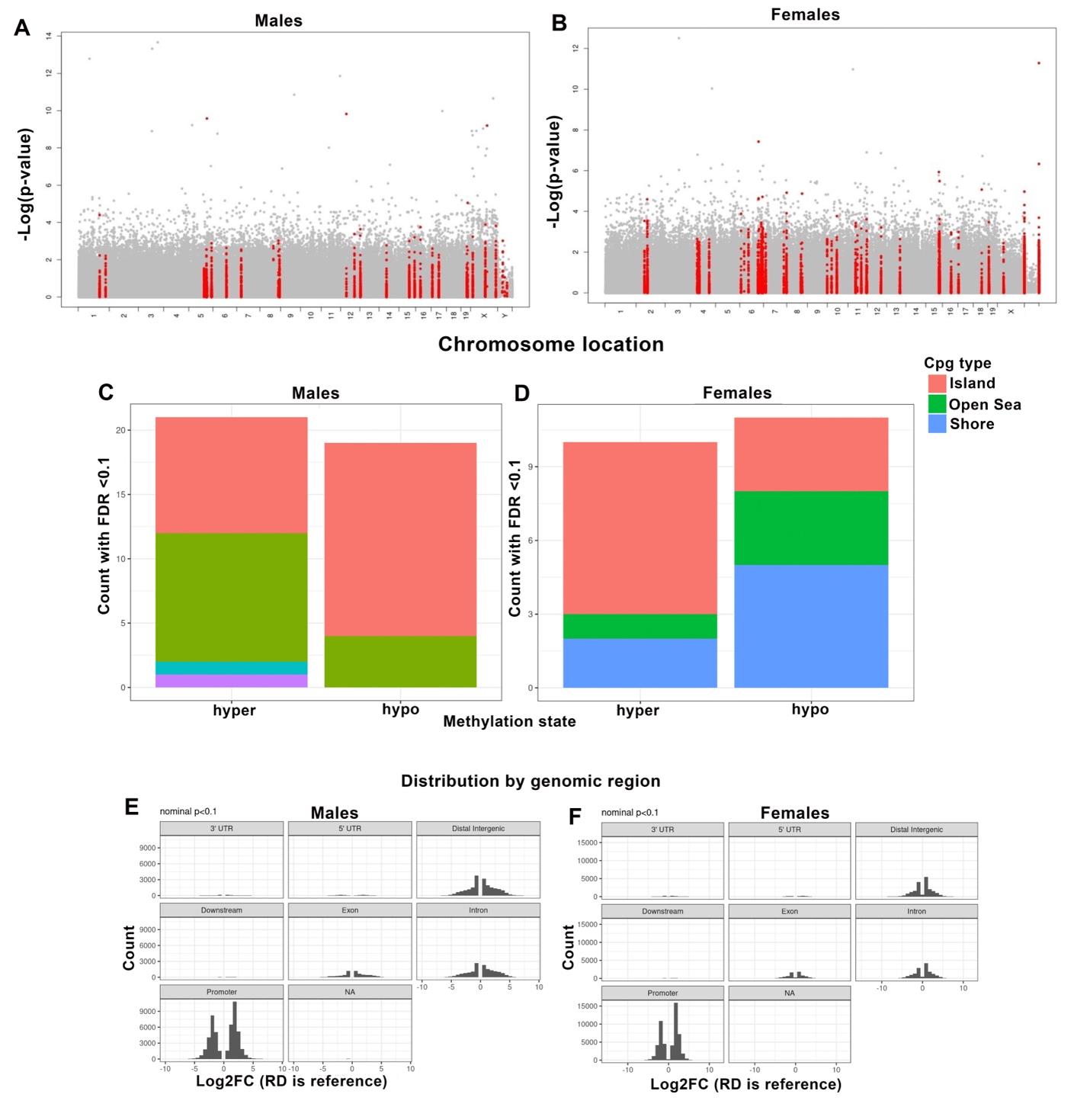

**Figure S6**. **A-B,** Manhattan plots for DNA methylation in 6-month-old male (**A**) and female (**B**) Off-HFD. Chromosomal position is depicted on the x-axis, - log(*p*-value) on the y-axis. The red line indicates nominal *p*-values<1x10^-14^. **C-D,** Stacked histogram of DNA methylation distribution in male (**C**) and female (**D**) Off-HFD showing the count of CpGs located in CpG islands (red), open sea (green), and shores (blue). **E-F**, Distribution of differentially methylated CpG sites in male (**E**) and female (**F**) Off-HFD.

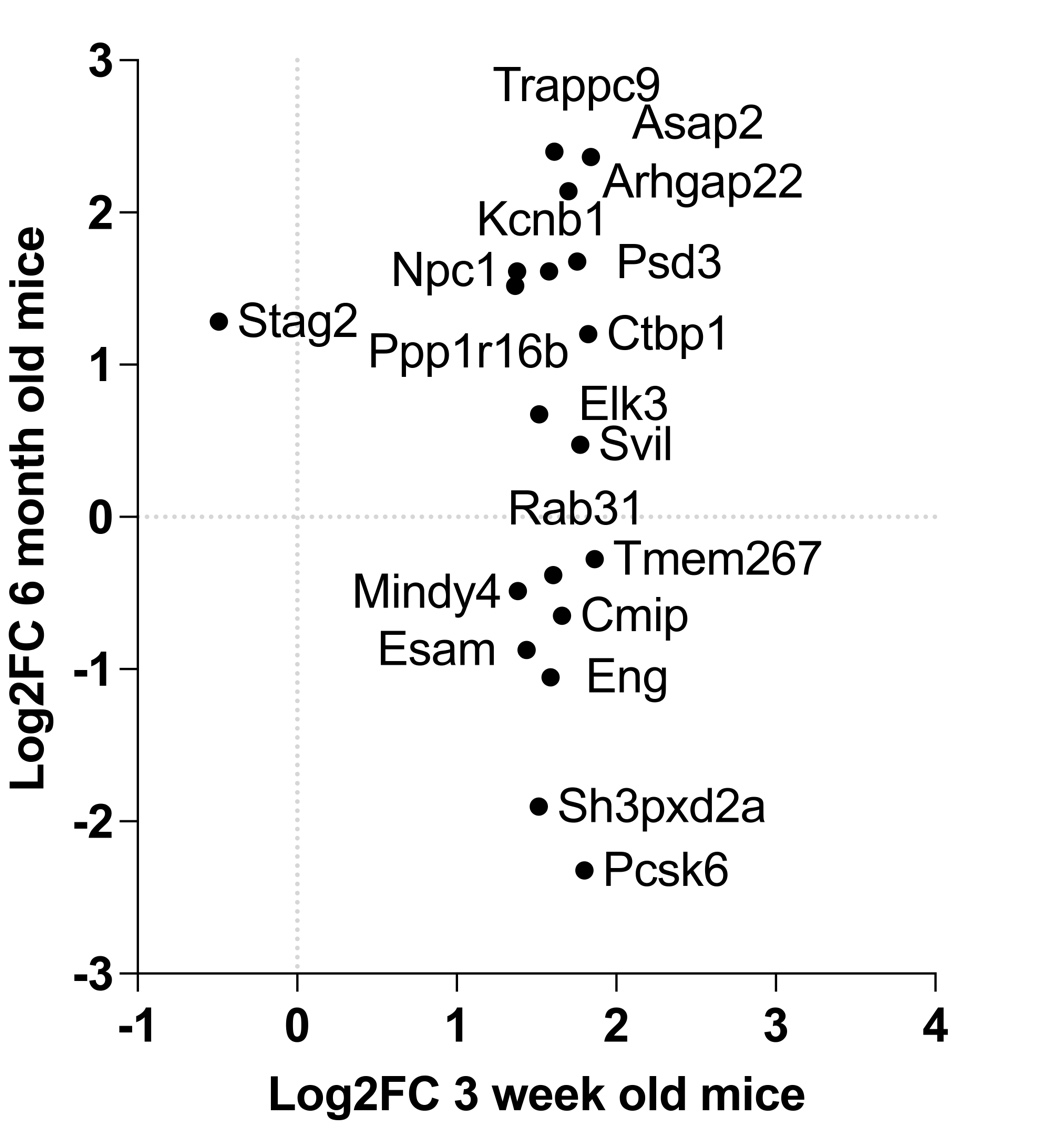

**Figure S7.** Comparison of RRBS data at individual CpG sites between 3-week-old and 6-month-old mice.

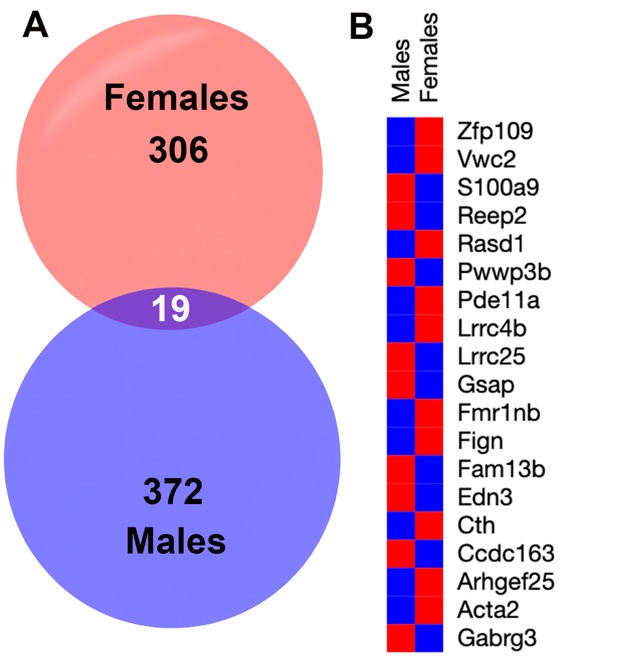

**Figure S8**. **Common differentially expressed genes (DEGs) affected by maternal obesity in male and female Off-HFD versus Off-RD mice.** **A,** Venn diagram showing the total number of DEGs identified in males and females, including 19 DEGs shared between sexes. **B,** Heatmap showing log2 fold-change values for the shared DEGs. All shared DEGs exhibited opposite sex-dependent patterns of regulation in response to maternal obesity.

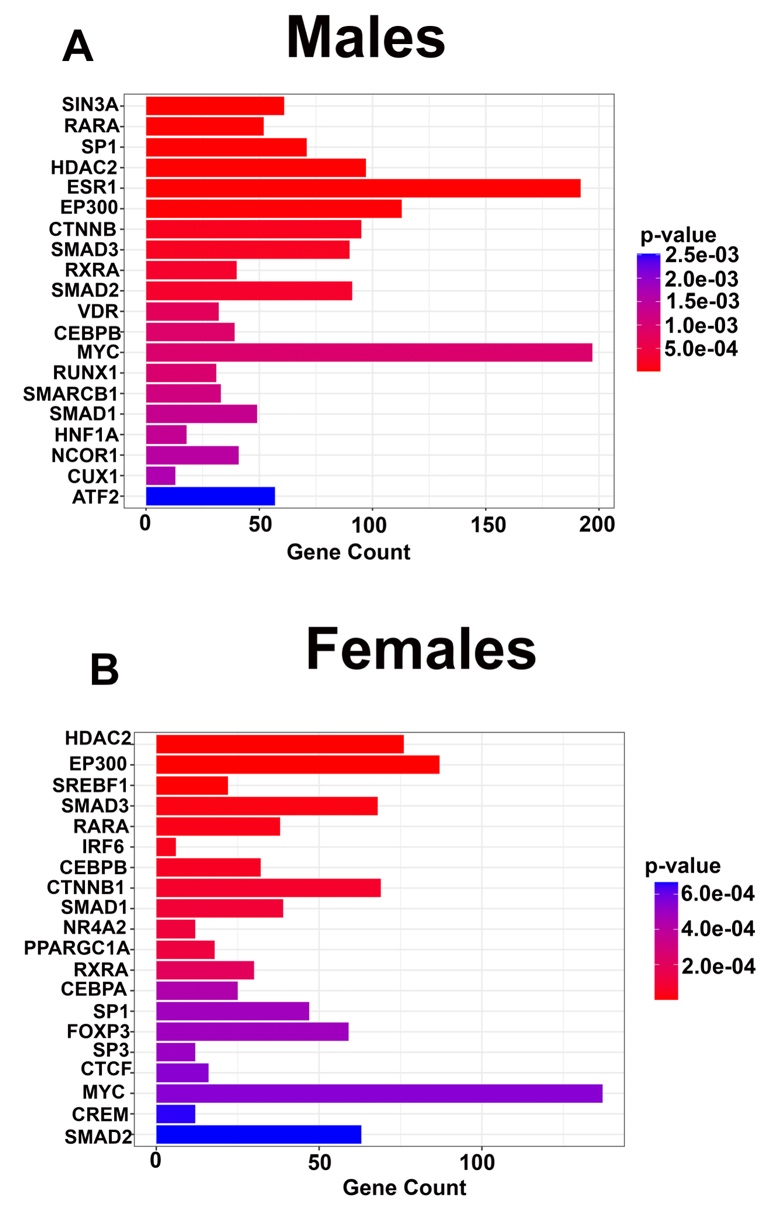

**Figure S9**. Significant modules from transcription factor protein–protein interaction (PPI) network enrichment analysis in male (**A**) and female (**B**) Off-HFD mice.
